# DDX41 is needed for pre-and post-natal hematopoietic stem cell differentiation in mice

**DOI:** 10.1101/2021.08.13.456151

**Authors:** Jing Ma, Nadim Mahmud, Maarten C. Bosland, Susan R. Ross

## Abstract

DDX41 is a tumor suppressor frequently mutated in human myeloid neoplasms. DDX41 binds to DNA/RNA hybrids and interacts with spliceosome component (1, 2). How it affects hematopoiesis is still unclear. Using a knockout mouse model, we demonstrate that DDX41 is required for mouse hematopoietic stem and progenitor cell (HSPC) survival and differentiation. Lack of DDX41 particularly affected myeloid progenitor development, starting at embryonic day 13.5. DDX41-deficient fetal liver and adult bone marrow (BM) cells were unable to rescue mice from lethal irradiation after transplantation. DDX41 knockout stem cells were also defective in ex vivo colony forming assays. RNASeq analysis of lineage-negative, cKit+Sca1+ cells isolated from fetal liver demonstrated that the expression of many genes associated with hematopoietic differentiation were altered in DDX41 knockout cells. Furthermore, altered splicing of genes involved in key biological processes were observed. Our data reveal a critical role for DDX41 in HSPC differentiation and myeloid progenitor development, likely through its regulation of gene expression programs and splicing.

**Significance:** DDX41 is a tumor suppressor in hematologic malignancies. However, whether DDX41 functions in hematopoiesis and myeloid cell differentiation is not known. Here we show that in mice, loss of DDX41 in hematopoietic stem cells (HSCs) leads to defects in hematopoietic development. The myeloid lineage was particularly affected as early as pre-natal stages. Transcriptional profiling of embryonic HSCs revealed that there were global changes in gene expression and splicing due to lack of DDX41. Collectively, the study uncovers a new function of DDX41 in HSC differentiation and could provide molecular targets for treatment of myeloid differentiation disorders.

## Introduction

Myelodysplastic syndrome and acute myeloid leukemia (MDS/AML) are diseases of familial and sporadic origin(3). DEAD-box helicase 41 (*DDX41*) is one of the most commonly mutated genes in both forms (1, 4–8). DDX41 belongs to a family of DEAD/H box RNA helicases whose members are involved in translation, ribosome biogenesis, nuclear-cytoplasmic transport, pre-mRNA splicing and nucleic acid sensing and pathogen recognition (7, 9–12). DDX41, which shows 99% amino acid identity in mice and humans, contains an N-terminal nuclear localization signal, the nucleic acid-interacting DEAD box, a helicase domain and a C-terminal Zn^2+^ finger (2, 13, 14). DDX41 is a tumor suppressor in MDS/AML; in familial disease, one mutated copy is inherited and somatic mutations in the second allele are found in leukemic cells.

It is thought that loss of DDX41 results in increased myeloid cell proliferation through its effects on splicing (1). Splicing factor (SF) mutations are common in MDS/AML (15). DDX41 interaction with spliceosomes and altered splicing may cause tumor suppressor inactivation or alterations in the balance of gene isoforms in MDS/AML (1). DDX41 is also a nucleic acid sensor (13, 16). It binds the first product of retroviral reverse transcription, RNA/DNA hybrids and then to Stimulator of Interferon Genes (STING), initiating an anti-viral response (2). DDX41 also senses RNA/DNA hybrids translocated from the mitochondria during influenza A infection (17). That DDX41 interacts with protein and RNA/DNA hybrids in viral nucleic acid sensing and with spliceosomes suggests an overlap in function.

We recently showed that *Ddx41* germline knockout in mice was embryonic lethal; to study its role in sensing of retroviruses, we used mice with a floxed allele (*Ddx41^f/f^*) and mice with macrophage and dendritic cell-specific gene deletion (2). The mice had normal lifespans, and aside from increased susceptibility to infection with murine leukemia virus, showed no immune cell defects or cancer. Thus, DDX41 loss in terminally-differentiated macrophages did not cause transformation and suggests that the protein acts at an earlier stage in AML/MDS.

Here, we show that HSC deletion of *Ddx41* causes defects in differentiation ex vivo and in vivo and depletion of myeloid lineages. Rather than leading to increased proliferation, as has been shown in other systems, *Ddx41* knockout in mouse HSCs decreased proliferation and differentiation (1, 18, 19). RNASeq analysis of DDX41-deficient fetal liver HSCs showed profound differences in expression of genes likely to play a role in differentiation and proliferation. Thus, HSC loss of DDX41 does not result in a pro-oncogenic phenotype, but inhibits their normal development.

## Results

### Mice lacking DDX41 in HSCs have hematopoietic defects

To create mice with DDX41-deficient HSCs, we generated VavCre^+^ *Ddx41^+/f^* males with one wild type and one floxed allele. We used VavCre^+^ males because this transgene is expressed in oocytes and causes germline deletion, confounding the studies (unpublished) (20). The mice were mated with *Ddx41^f/f^* females to generate VavCre+ *Ddx41^-/-^* knockout (KO), VavCre+ *Ddx41^+/-^* (het), and wild type (WT) *Ddx41^+/f^* and *Ddx41^f/f^* offspring. KO mice were born at the predicted Mendelian frequency (Fig. 1A) and appeared normal at birth (*SI Appendix,* Fig. S1A). At postnatal day 2-3 (PD3), the KO mice demonstrated failure to thrive – their growth was stunted in comparison to their WT or het littermates and they appeared anermic (*SI Appendix*, Fig. S1A). Most KO mice did not survive beyond day 15 (Fig. 1B). At PD5, the major organs of KO mice, particularly the spleen, were reduced in size and there were fewer erythroblasts and erythrocytes in the liver, spleen and BM (*SI Appendix*, Fig. S1A, S1B). Peripheral blood smears indicated anemia and complete blood count (CBC) analysis showed decreased leukocytes, erythrocytes and platelets in PD3 KO mice (*SI Appendix*, Fig. S1B, S1C). No differences were seen between WT and het mice. Because of this lack of a difference, as well as in gene expression analysis (see below), WT and het mice were used interchangeably in this study.

**Fig. 1.**
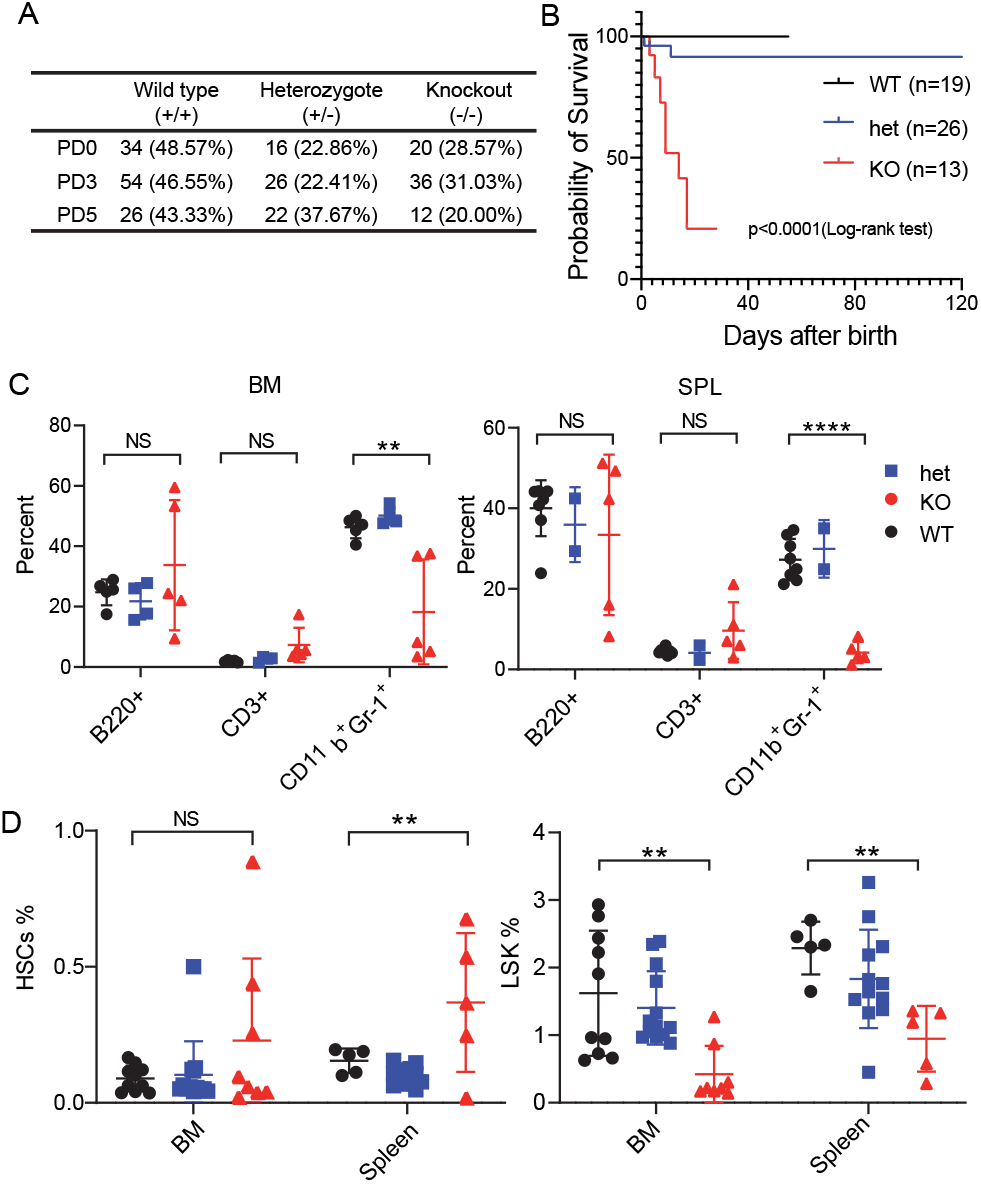
DDX41 KO mice fail to survive postnatally. A) DDX41 KO mice were born with appropriate Mendelian frequency. B) Postnatal survival of DDX41 KO mice. P values were measured by the Log-rank test. C) Frequency of hematopoietic cells in bone marrow (BM) and spleen (SPL) at PD3. BM: WT, n=5; het, n=4; KO, n=5. SPL: WT, n=8; het, n=4; KO, n=7. D) Frequency of HSPCs (LSK+) decreased in KO while HSCs (LSK/CD150+/CD48-) were comparable between WT and KO PD3 mice. BM: WT, n=10; het, n=13; KO, n=8. SPL: WT, n=5; het, n=12; KO, n=5. One-way ANOVA with post-hoc Tukey test was used to determine significance: *, *P*≤0.02, **, *P*≤0.01; ***, *P*≤0.001; ****, *P*≤ 0.0001; ns, not significant. For all experiments, mice were derived from a minimum of 3 litters.

We analyzed the effect of DDX41 loss on B cells, T cells and myeloid cells in PD3 BM and spleen. We observed decreased cell numbers in all three populations in KO spleens (*SI Appendix*, Fig. S1D). However, the B and T cell percentage was not different between genotypes, while the myeloid cell percentage significantly decreased in the KO BM and spleens compared to WT and hets (Fig. 1C). Similar results were seen at PD5 (*SI Appendix*, Fig. S1E).

We also tested whether HSCs were affected by DDX41 deficiency. During development, HSCs colonize the fetal liver, followed by translocation to neonatal BM and spleen (21). At PD3, the HSC percentage was the same or slightly increased in the KO BM and spleens compared to WT and hets (Fig. 1D, *SI Appendix*, Figs. S1F and S2B). However, the LSK^+^ population frequency and numbers were significantly decreased (Fig. 1D and *SI Appendix*, Fig. S1F). These results show that lack of DDX41 affects hematopoietic cell development in general and differentiation of myeloid lineage cells.

### Defects in HSC differentiation in DDX41 KO embryos

HSCs expand in the fetal liver from embryonic day 12.5-14.5 (22). We next examined the fetal liver stem cell population. Similar to neonatal mice, the HSC frequency was not affected at E13.5 or E14.5 (Fig. 2A; *SI Appendix*, Figs. S2A, S2B). However, the KO fetal liver LSK+ population was significantly reduced at E14.5 but not E13.5, compared to WT or hets (Fig. 2A; *SI Appendix*, Figs. S2A, S2B).

**Fig. 2.**
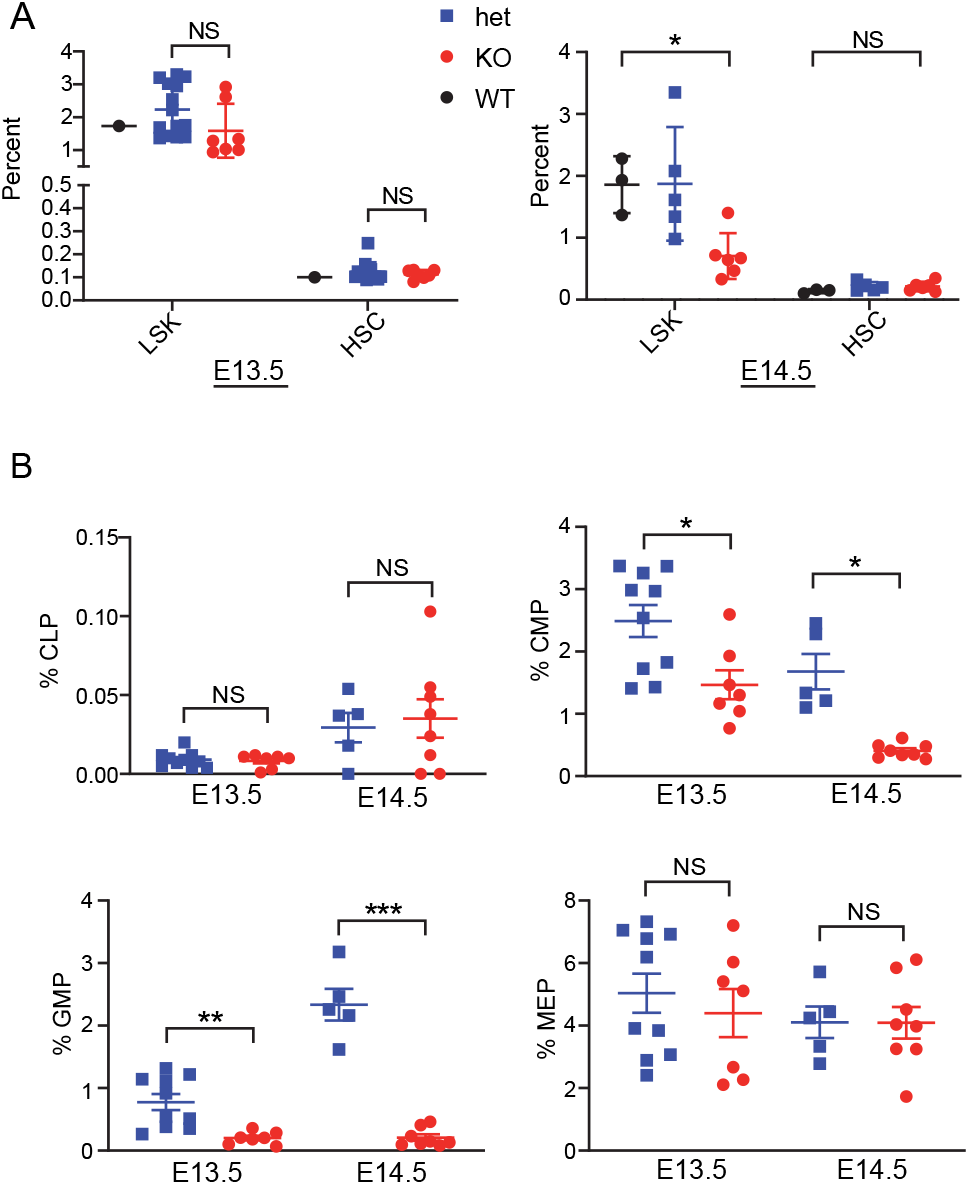
HSC defects begin prenatally in DDX41 KO mice. A) Cells isolated from prenatal liver were examined for hematopoietic stem/progenitor cells (LSK+) or hematopoietic stem cells (HSCs) at day E13.5 (KO, n=7; het, n=14; WT, n=1) and E14.5 (KO, n=6; het, n=5; WT, n=3). The mice were derived from 3 litters for each experiment. Unpaired two-tailed t-test was used for E13.5 and one-way ANOVA with post-hoc Tukey test was used for E14.5 to determine significance: *, *P*≤<0.05. B) Cells were also examined for lineage-committed progenitor markers (CMP: Lin^-^/c-Kit^+^/Sca-1^-^/CD34^+^/CD16/32^-^; GMP: Lin^-^/c-Kit^+^/Sca-1^-^/CD34^+^/CD16/32+; MEP: Lin^-^/c-Kit^+^/Sca-1^-^/CD34^-^/CD16/32^-^; CLP: Lin^-^/CD127^+^/c-Kit^low^/Sca-1^low^). E13.5: het, n=10; KO n=7. E14.5: het, n=5; KO, n=8. Unpaired two-tailed t-tests were used to determine significance: *, *P*≤0.05; **, *P*≤0.01; ***, *P*≤0.001. The mice were derived from 3 litters for each experiment.

We further investigated lineage-committed progenitors from E13.5 and E14.5 KO and het fetal livers (Fig. 2B, *SI Appendix*, Fig. S2C). A lower percentage of granulocyte-monocyte progenitors (GMPs) and common myeloid progenitors (CMPs) in KO vs. het livers was detected at both time points. Neither the percentages of common lymphoid progenitors (CLPs) or megakaryocyte–erythroid progenitors (MEPs) from KO fetal liver showed differences compared to hets. These data show that while all LSK+-derived lineages were diminished, the myeloid lineage was most affected by DDX41 loss.

### DDX41 depleted HSCs fail to expand ex vivo and in vivo

We also tested whether DDX41 deficiency affected HSC ex vivo differentiation and BM repopulation. In a pilot CFU assay, we found very few colonies when 20,000 KO E14.5 fetal liver cells were plated (not shown). To obtain sufficient numbers of KO colonies, 5-fold more KO than het cells were plated. Nonetheless, significantly fewer BFU-E, CFU-GM, and CFU-GEMM colonies were obtained with KO vs het fetal liver cells (Fig. 3A). Moreover, the CFU-GM were smaller in area, suggesting a slower proliferative rate (*SI Appendix*, Figs. S3A, S3B).

**Fig. 3.**
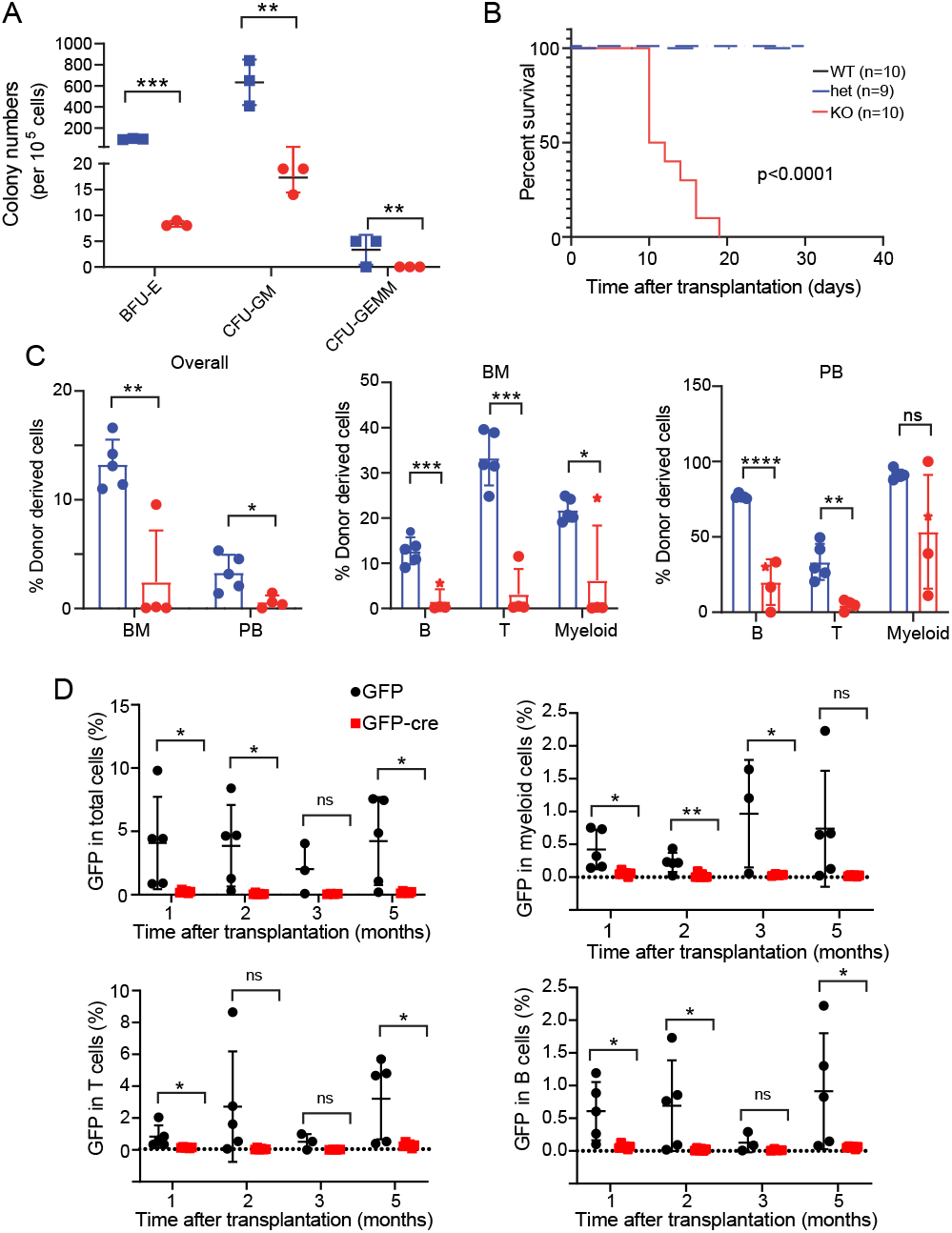
HSCs from DDX41 KO embryos and adult bone marrow have compromised repopulating capacity. A) E14.5 fetal liver cells from KO, het and WT mice were seeded for CFU assays. Data were collected from three independent experiments. B) Transplantation of DDX41 KO and het E14.5 fetal liver cells into lethally irradiated recipients. Kaplan Meier Curve was compared using Log-rank test. C) Left panel: percentage of donor derived cells in BM and peripheral blood (PB) 17 days after transplantation with E14.5 fetal liver cells. Percentage of donor-derived B cells, T cells and myeloid cells in BM (middle) and PB (right). N≥4 in each group of mice. D) Transplantation of DDX41-deficient adult bone marrow cells into lethally-irradiated recipients. Percentage of donor-derived total cells, T cells, B cells, and myeloid cells in PB up to 5 months after the transplantation. N>3 for each group of mice. Unpaired two-tailed t-tests were used to determine significance: *, *P*≤0.05, **; *P*≤0.01; ***, *P*≤0.001; ****, *P*≤0.0001.

Next, we tested whether E14.5 fetal liver cells repopulated the BM of lethally-irradiated recipients. First, we transplanted 500,000 WT, het and KO CD45.2^+^ fetal liver cells into CD45.1^+^ recipients. All recipients that received KO cells died between 10-20 days after transplantation, indicating that KO HSCs had limited engraftment ability (Fig. 3B). We repeated the transplantation experiment by injecting 20 times more fetal liver cells (1 × 10^7^) from KO and het embryos. Four of five KO HSC recipients showed failure to thrive by 17-days post-transplantation, while all recipients of WT and het HSCs demonstrated recovery, as determined by weight and CBCs (*SI Appendix*, Figs. S3C, S3D).

Because the majority of the KO mice were moribund, all mice were euthanized at day 17, and BM and peripheral blood were analyzed for donor cells. There were fewer donor-derived cells in the BM and blood of mice that received KO HSCs and all 3 lineages (B, T and myeloid) were decreased (Fig. 3C). The failure to repopulate was not a homing defect, since at 16 hr post-transplantation, equal percentages of WT and KO LSK+ cells were recovered from BM (*SI Appendix*, Fig. S3E).

Because other studies showed that DDX41 knockdown in adult HSCs increased proliferation, we tested whether DDX41 KO HSCs isolated from adult BM repopulated lethally-irradiated mice. Lineage-negative cells from 6-12 week old DDX41^fl/fl^ mice were transduced with a MSCV vector expressing Cre-recombinase and GFP, or GFP alone; Cre expression causes the loss of exons 7 - 9 (*SI Appendix*, Fig. S4A). The knockout allele was clearly detected by PCR in cells transduced with the Cre-containing vector (*SI Appendix*, Fig. 4B). FACS analysis showed that 30% of LSK+ cells were GFP+ at 48 hr post-transduction (*SI Appendix*, Fig. S4C). These cells were transplanted into lethally-irradiated recipients. GFP-transduced cells were detectable in transplanted mice for up to 5 months after transplantation in all populations (B, T and myeloid), while recipients that received Cre-transduced cells were gone in all populations by 1 month after transplantation (Fig. 3D).

**Fig. 4.**
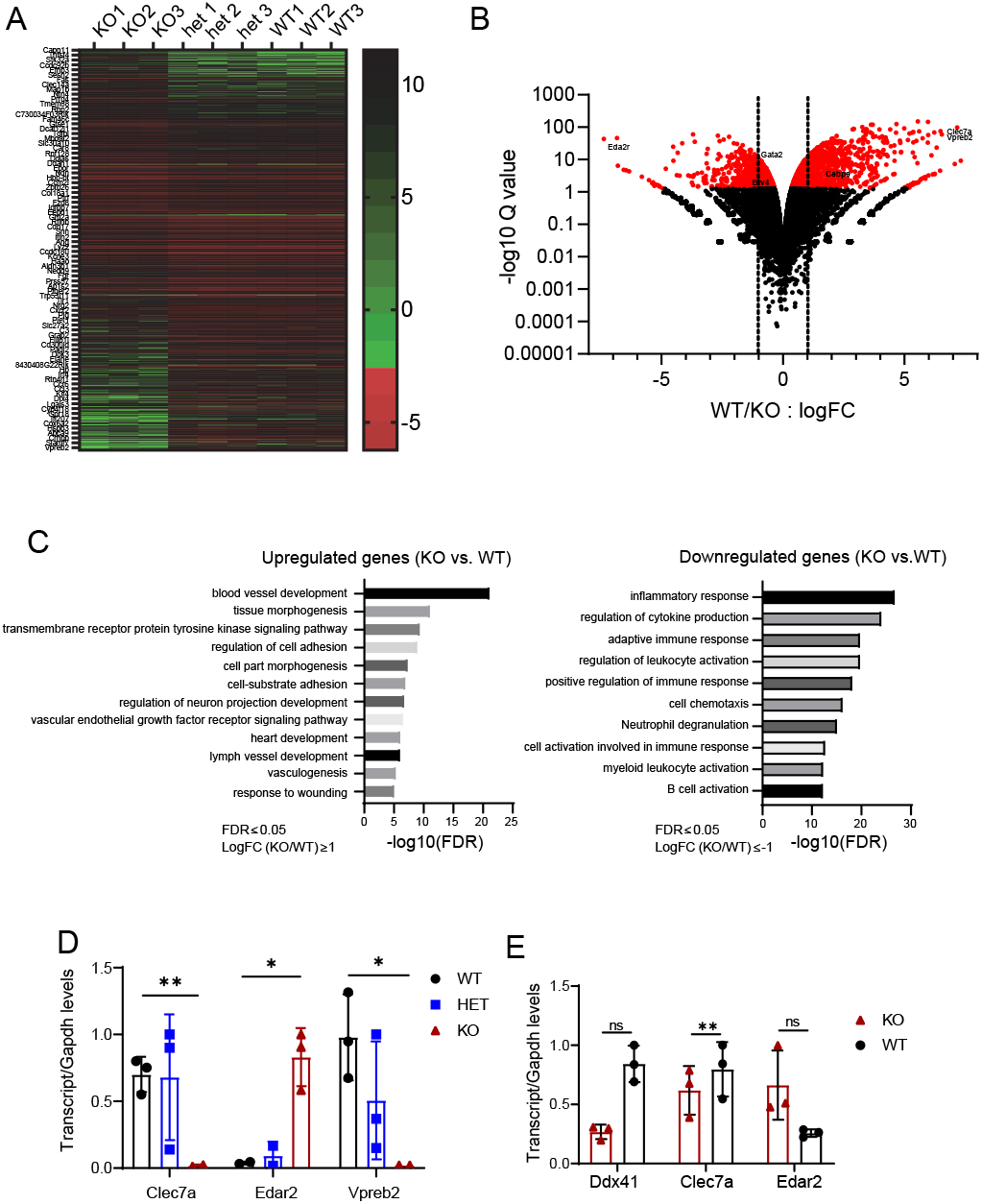
Loss of DDX41 is linked to global alternations in gene expression. A. Heatmap representation of genes differentially expressed between WT, het and KO. logFC ≥ 1 or ≤ -1. FDR ≤ 0.05. B. Volcano plot displaying differentially expressed genes between WT and KO, Cut-off: logFC ≥ 1 or ≤ -1, FDR ≤ 0.01. Pseudogenes were excluded. Red points represent genes above the FDR threshold. Vertical lines separate genes by denote genes with 2-fold or greater difference. N=3 replicates per condition. C. Gene ontology enrichment analysis of genes higher (logFC ≥ 1) or lower represented (logFC ≤ -1) in KO vs. WT. FDR ≤0.05. D. Validation of differentially expressed genes in WT, het and KO LSK+ cells by quantitative real-time PCR. Expression levels were normalized to Gapdh. E. PCR validation of differentially expressed genes in WT, het and KO BMDMs. Significance was evaluated by unpaired two-tailed t-test between WT and KO. *, *P*≤0.05; **, *P*≤0.01. N=3 replicates per experiments.

### Loss of DDX41 in LSK+ cells results in gene expression alterations

We next performed whole RNAseq of LSK+ cells isolated from WT, het and KO E14.5 livers (three mice of each genotype); 22320 annotated genes were analyzed. The KO mice showed significantly different gene expression levels than WT or het mice, which were similar to each other (Fig. 4A; *SI Appendix*, Fig. S5A). DDX41 deficiency resulted in 415 upregulated and 682 downregulated genes in KO vs WT/het cells (2-fold or greater) (Fig. 4B; *SI Appendix*, Dataset S1). GO term enrichment analysis of the genes more highly-expressed in WT/het compared to KO LSK+ cells revealed signatures of inflammation, cytokine production and adaptive immune response (top 3 categories), myeloid and leukocyte differentiation and activation, and neutrophil degranulation (Fig. 4C). In contrast, genes involved in blood vessel development and vasculogenesis were more highly represented in KO than WT/het LSK+ cells (Fig. 4C). Since HSCs are believed to arise from primitive endothelium, this suggest that lack of DDX41 arrests cells at an early step in the differentiation pathway.

We also found that differentially expressed transcription factors (TFs) in KO and WT/het cells were enriched in GO terms for embryonic development and cell differentiation (*SI Appendix*, Fig. S5B). Factors thought be important in HSC differentiation, such as GATA2 and ETV4, were slightly higher in KO cells, while CEBPE, believed to play a role in granulocyte progenitor cells, was higher in WT/het cells (Fig. 4B) (23–25). Similarly, several cell surface receptors were differentially expressed in KO and WT/het cells, with down-regulation of receptors involved in cytokine signaling, inflammatory responses and leukocyte activation in the KO LSK+ population (*SI Appendix*, Fig. S5C).

We used RT-qPCR to validate the RNAseq data in E14.5 LSK+ cells. Vpreb2 and Clec7a, genes involved in immune response, were more highly expressed in WT/het LSK+ cells, while Edar2, involved in ectodermal tissue development, was more highly expressed in KO cells (Fig. 4D). There were also differences in Clec7 and Eda2r expression in WT and DDX41 KO BMDMs isolated from LyCreDDX^fl/fl^ mice(2) (Fig. 4E); Vpreb2 is not expressed in macrophages. This suggests that DDX41 also regulates gene expression in macrophages, but that expression of these genes is sufficient for differentiated macrophage but not HSC differentiation.

### Loss of DDX41 alters splicing

DDX41 regulates splicing in human cell lines, C. elegans and zebrafish embryos (1, 18, 26, 27). We analyzed alternative splicing using rMATS (*SI Appendix*, Datasets S2-S6) (28). There were 367 differential splicing events between WT and KO. Skipped exons (SE) and retained introns (RI) were more frequently altered than alternative 5’ splice site (A5SS), alternative 3’ splice site (A3SS) or mutually exclusive exons (MXE) in KO vs. WT LSK+ cells (Fig. 5A). GO term analysis of altered gene splicing events (n=350, FDR < 0.05, inclusion difference > 10%) revealed signatures of chromatin organization, DNA repair, mRNA processing and cellular amide metabolism (Fig. 5A). Genes with increased SEs were enriched for terms related to RNA splicing, mRNA transport, and post-translational modification (n=175, FDR < 0.05, inclusion difference > 10%), while those with elevated RIs in KO cells clustered in the GO terms of ribosome biogenesis, DNA damage response, protein modification and mRNA processing (n=106, FDR < 0.05, inclusion difference > 10%) (Figs. 5B, 5C). Differential splicing was also found in genes encoding TFs, SFs and receptors (*SI Appendix*, Dataset S7).

**Fig. 5.**
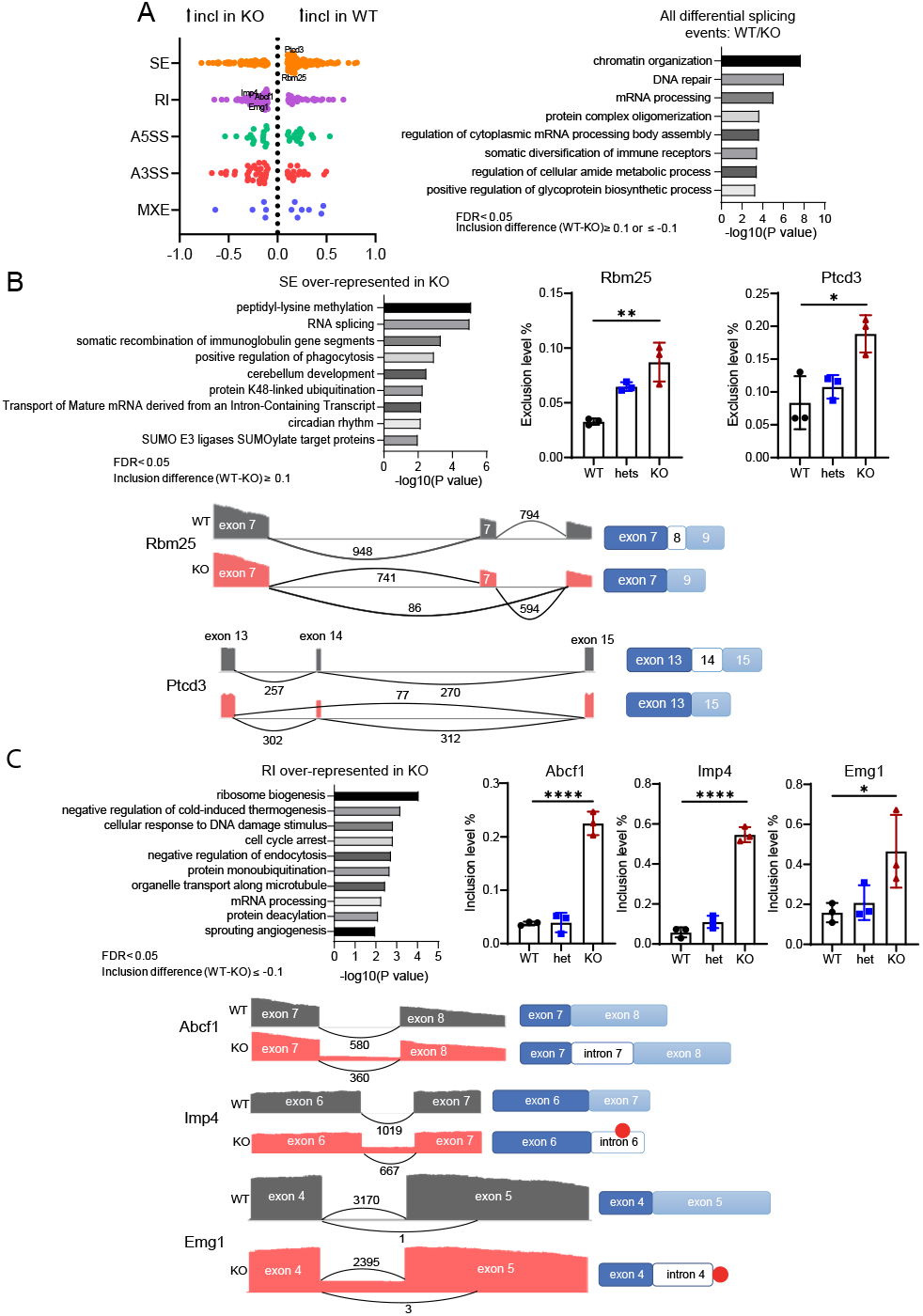
Loss of DDX41 led to alternative splicing. A. Distribution of inclusion difference for major types of alternative splicing events (left). FDR < 0.05, inclusion difference> 10%. Types of alternative splicing events: Skipped exon (SE), alternative 5’ splice site (A5SS), alternative 3’ splice site (A3SS), mutually exclusive exons (MXE), retained intron (RI). GO analysis of all differential splicing events between WT and KO (right). B. GO analysis of over-represented SE in KO/WT (upper left). Splicing difference between WT, het and KO was shown by exclusion level (the percentage of spliced-out product in both spliced-in and spliced-out products) (upper right). Visualization of skipped exon by Sashimi Plot for Rbm25 and Ptcd3 in KO LSK+ cells compared to WT LSK+ cells (lower). Significance was calculated by one-way ANOVA with post-hoc Tukey test. *, *P*≤0.05; **, *P≤0.01.* N=3 replicates per experiment. C. GO analysis of over-represented RI in KO and WT cells. Graph representing splicing difference by inclusion level (the percentage of spliced-in product) between WT, het and KO cells. Image Lab software (Bio-Rad) was used for gel quantification. Band intensity was normalized to Gapdh. Visualization of retained intron by Sashimi Plot for Abcf1, Imp4, Emg1 between WT and KO. Significance was evaluated by one-way ANOVA with post-hoc Tukey test. *, *P*≤0.05; ****, *P*≤0.0001. N=3 replicates per experiment.

Candidates involved in mRNA processing (Rbm25) and ribosome biogenesis and translation (Abcf1, Imp4, Emg1, Ptcd3) were validated by RT-qPCR (Figs. 5B, 5C; *SI Appendix*, Figs. S6, S7A and S7B). Splicing of these genes was similar in WT and hets (*SI Appendix*, Figs. S6 and S7). DDX41 also affected splicing in differentiated DDX41-deficient BMDMs, with elevated RIs in Abcf1 and Imp4 and increased SEs in Ptcd3 and Rbm25; however, no altered splicing of Emg1 was observed (*SI Appendix*, Figs. S8A and S8B).

DDX41-deficiency resulted in retention of intron 7 in Abcf1, intron 6 in Imp4 and intron 4 in Emg1 and skipping of exon 8 in Rbm25 and exon 14 in Ptcd3. For Emg1 and Imp4, the retained intron identified in KO LSK+ RNAs results in a premature termination codon, likely leading to transcript nonsense-mediated decay (NMD), while the Abcf1 retained intron added additional in-frame coding sequences. The altered splicing of Ptcd3 and Rbm25 resulted in in-frame proteins with exon loss (Figs. 5B, 5C; *SI Appendix*, Figs. S6 and S7). Thus, altered splicing likely affected function of proteins important to many pathways.

## Discussion

DDX41 is a nucleic sensor that binds DNA and DNA/RNA hybrids generated during virus infection (2, 13, 16, 17). Here we show using a knockout mouse model that DDX41 plays a critical role in mouse HSC survival and development, and particularly disrupts the differentiation of myeloid lineage cells. Loss of DDX41 in fetal LSK+ cells also led to global changes in gene expression and altered splicing, suggesting that DDX41 acts as a regulator of hematopoietic development.

DDX41 knockout had an effect as early as day E14.5, when the frequency of fetal liver LSK+ cells underwent a severe loss while that of HSCs remained stable. Given that HSCs constitute a subset of the LSK+ population, this suggests that DDX41 is required for differentiation of a progenitor population downstream of HSCs. Recent studies using single-cell RNAseq of adult BM HSCs have indicated that this progenitor population is heterogeneous (29–32). DDX41 may be required for differentiation of a subset of the progenitor population, currently undefined by marker analysis. This could explain why lack of DDX41 led in particular to decreases in CMPs as early as day E13.5.

Although a decreased percentage of CMP and GMP progenitors occurred at E13.5, the myeloid lineage was not the only subset affected by lack of DDX41. While lymphoid and erythroid cell percentages were not affected in DDX41 KO newborns, the absolute numbers of both subsets were decreased; the deficit in the erythroid lineage led to severe anemia, which was the likely cause of death (*SI Appendix*, Fig. S1B). DDX41 plays a role in the maintenance and proliferation of HSCs and HSPCs, since DDX41-deficient cells were defective in both colony assays and transplantation experiments. Given that CMPs give rise to myeloid and erythroid cells, it was unexpected that progenitor MEPs were not affected by prenatal DDX41 loss (Fig. 6A). In a parallel lineage branching model, it was proposed that heterogeneous CMPs yield only erythrocytes or myeloid cells after transplantation, and that the divergence of the two lineages develops within progenitors upstream of CMP and GMP (33). DDX41 may thus be especially important for a subset of LSK/CMP cells that go on to contribute to the myeloid lineage. What is not known is how far back in the lineage DDX41 activity distinguishes the progenitor subset.

**Fig. 6.**
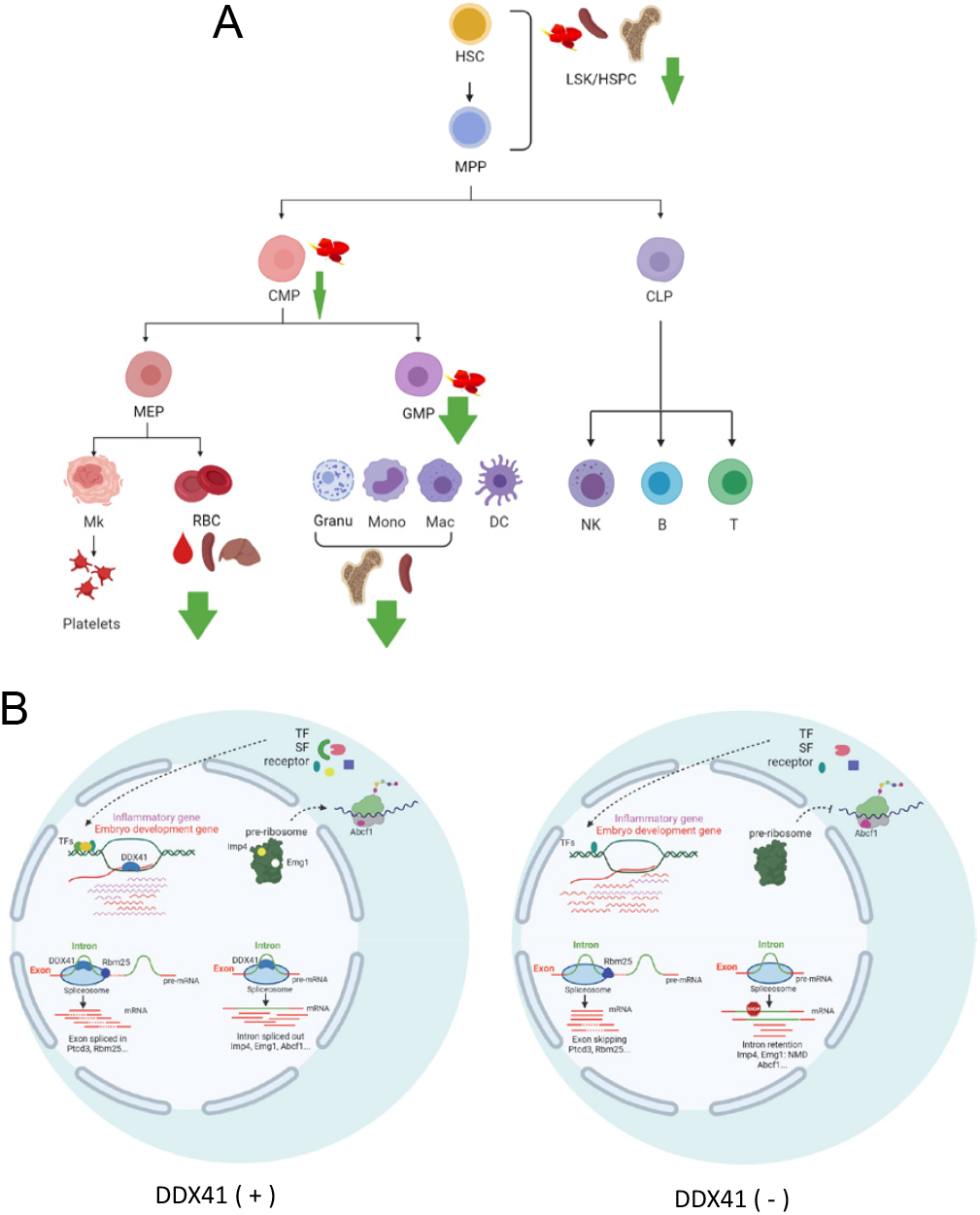
DDX41 causes defects in the differentiation of mouse HSCs. A. Overview of mouse HSC differentiation in DDX41-defcient mice. The thickness of arrow indicates greater extent of the effect. Tissues examined are shown beside the cells. LSK+ cells, which includes HSCs and MPPs, decreased both prenatally and postnatally, while HSCs are not affected by the absence of DDX41. A reduction was seen in fetal liver CMPs and a greater decrease in GMPs and myeloid cells. RBCs were decreased in peripheral blood, neonatal spleen and liver and severe anemia was observed. B. Model of DDX41-mediated gene regulation in mouse HSPCs. DDX41 positively and negatively regulates expression of genes involved in inflammation and embryo development, respectively. DDX41 is also required for appropriate splicing of critical genes associated with ribosome biogenesis, translation initiation and mRNA processing. Alternative splicing in DDX41-deficient cells leads to decreased protein production through NMD (Imp4, Emg1), changes in isoforms and aberrant production of functional regulators (TF, transcription factor; SF, splicing factor; receptor), which likely in turn affect gene transcription.

Inflammation is critical for HSPC proliferation and differentiation through increased expression of adhesion and hematopoietic growth factors (34, 35). We showed that loss of DDX41 in fetal liver cells impedes mouse HSPC survival and normal development, and whole transcriptome profiling of DDX41-deficient LSK+ cells showed decreased expression of genes involved in inflammatory and immune responses which are likely required for HSPC differentiation. Recently, it was reported that DDX41 regulates homeostasis by suppressing the cGAS-STING inflammatory pathway in zebrafish and human HSPCs, which is the opposite of what we observed here (18, 19). Although HSC self-renewal and differentiation is highly conserved among vertebrates, zebrafish and mouse have different hematopoiesis timelines and erythropoiesis occurs at distinct sites (36–38). Thus, it is possible that DDX41 functions distinctly in different species.

Whole transcriptome profiling in different species shows that DDX41 functions in RNA splicing (1, 18, 26, 27). While our RNASeq data showed that DDX41 had the greatest effects on transcript levels in LSK+ cells, the data also showed that DDX41 loss altered exon skipping and intron retention. In contrast to our results, in HeLa cells A3SS, A5SS and SE were all regulated by DDX41, while in *C. elegans* upon gene deletion, A3SS represented the predominant splicing defect (26, 27). Whether these differences are species-or cell type-specific remains to be determined.

DDX41 mutations occur in familial and sporadic AML/MDS (1, 6, 39). In silico analysis of transcripts expressed in AML/MDS and DDX41 pull-down experiments suggested that is involved in splicing and aberrant regulation of critical growth regulatory genes (1). Moreover, DDX41-depleted human adult CD34+ cells exhibited more proliferation and greater colony formation, supporting its role in oncogenesis (1). Instead, we found that lack of DDX41 resulted in lower cell proliferation, loss of LSK+ and fully differentiated hematopoietic cells and an inability to repopulate the BM. This suggests that in mouse fetal HSPCs, DDX41 plays a different role. Indeed, many of the genes dysregulated in LSK+ cells showed similar expression and splicing patterns when DDX41 is depleted from macrophages, but did not affect their function.

We suggest that DDX41 regulates HSPC survival and development through its participation in transcription, perhaps by binding DNA/RNA hybrids and TFs (Fig. 6B). Simultaneously, DDX41 acts as part of splicing machinery, modulating alternative splicing leading to NMD and altered ratios of isoforms crucial to key biological processes like ribosome biogenesis, translation initiation and mRNA processing. Finally, the affected TF, SF and receptor translation products in turn modulate transcription and contribute to the DDX41 KO phenotype.

## Methods

### Mice and genotyping

Mice were bred at the University of Illinois at Chicago (UIC). DDX41^fl/fl^ mice (C57BL/6N) were constructed by TaconicArtemis GmbH and were previously described, as were LyCreDDX^fl/fl^ mice (2). VavCre (B6.Cg-Commd10Tg(Vav1-icre)A2Kio/J) mice were a gift from Kostandin Pajcini (40). C57BL/6N and B6.SJL-Ptprca Pepcb/BoyJ (CD45.1) mice were purchased from Jackson Laboratory. Both male and female mice were used for experiments, except the BM transplantation studies. All mice were housed according to the policies of the UIC Animal Care Committee (ACC); all studies were performed in accordance with the recommendations in the Guide for the Care and Use of Laboratory Animals of the National Institutes of Health. The experiments were approved by the UIC ACC (protocol no. 18-168). Primer sequences for genotyping are in *SI Appendix*, Table S1.

### Timed mating

Timed mating of female DDX41^fl/fl^ mice and male VavCre+ DDX41^fl/f+^ mice were carried out. Pregnant females were euthanized at day E13.5 and E14.5 and fetal liver cells were harvested for subsequent analyses.

### BM and splenocyte isolation

Tibiae and femurs were dissected and cells were collected by flushing. Livers and spleens were homogenized, red blood cells were lysed with ACK buffer, tissues were homogenized and filtered through a 70 μm cell strainer. Single cell suspensions were prepared in 1% fetal bovine serum (FBS)-PBS.

### Flow cytometry and sorting

Cells were incubated in blocking solution (Fc-block in 1% FBS-PBS) and then stained with fluorescently-labeled antibodies. LSK+ cells were identified by mouse lineage antibody cocktail (B220-FITC, CD3-FITC, CD11b-FITC, Gr-1-FITC, Ter119-FITC), ckit-APC and Sca-1-PE. HSCs were identified by CD150-PE/Cy7, CD48-APC/Cy7 and LSK. CD16/32-APC/Cy7, CD34-PE/Cy7, LSK, CD127-PerCp/Cy5.5 were used for analysis of lineage-committed progenitors (CMP, GMP, MEP, CLP)(41). CD11b-FITC and Gr-1-PE were used to stain myeloid cells. Antibody information can be found in *SI Appendix*, Table S2. DAPI was added to exclude dead cells. Flow cytometry experiments were performed on a BD LSR Fortessa with HTS and analyzed using FlowJo software. A Beckman MoFlo Astrios Cell Sorter was used for cell sorting. 15,000 to 20,000 LSK+ cells were sorted; the purity of the population was > 98%.

### BM rescue

Female recipient B6.SJL-Ptprca Pepcb/BoyJ mice were lethally irradiated (8.5 Gray; L Shepherd Model 143-68 Cesium 137 Irradiator) one day before transplantation. E14.5 fetal liver cells were harvested and single cell suspensions prepared. Cells from 2-3 mice were pooled for each genotype and 1 × 10^6^ cells were injected retro-orbitally into each recipient, or as indicated. Survival was monitored for up to 30 days.

### MSCV transduction

Retroviral vectors were produced in 293T cells, using a three-plasmid system (MSCV-Cre-GFP (Addgene plasmid #24064), MLV Gag-Pol/pHit111 and MLV Env/pHIT60) (42). Lineage depletion of fetal liver cells was completed following the manufacturer’s protocol (Miltenyi, 130-090-858). After culturing in stem cell media (SFEM media with mTPO, IL-3, IL-6, SCF), the cells were transduced with the MSCV-Cre-containing retrovirus. Reagent information is found in *SI Appendix*, Table S2.

### Homing assays

E14.5 liver cells were harvested from WT, het or KO mice (CD45.2). After lineage depletion, 5 × 10^5^ lineage-negative cells were injected retro-orbitally into lethally-irradiated B6.SJL-Ptprca Pepcb/BoyJ mice. Fetal liver cells were analyzed by flow cytometry for the number of injected LSK+ cells. At 16 h after transplantation, recipient BM cells were analyzed by flow cytometry. The absolute number of donor-derived cells in BM was determined and the percentage of homed cells/injected cells was calculated.

### Colony-forming unit assays

The protocol “Mouse Colony-Forming Unit (CFU) Assays using MethoCultTM” (Stemcell Technologies Canada Inc.) was used. Fetal liver cells were harvested from E14.5 embryos. Red blood cells were lysed and single cell suspensions were prepared in Iscove’s Modified Dulbecco’s Medium (IMDM) with 2% FBS. 20,000 to 100,000 cells were seeded in 35 mm culture dishes in duplicate culture experiments. Burst-forming unit-erythroid (BFU-E), granulocyte or macrophage progenitor (CFU-GM), and multi-potential progenitor cells (CFU-GEMM) were identified by their morphology 12-14 days after seeding. Colonies were captured with a Keyence BZ-X710 microscope. Colony number and size were measured by BZ-X Analyzing software.

### RNA library construction and sequencing

RNA was isolated from 1.5 × 10^4^ E14.5 fetal liver LSK+ cells from WT, het and KO mice (3 each) by RNeasy Micro Kit with DNase I (Qiagen). Quality control was done by Qubit RNA assay. Library construction and sequencing were performed by Novogene (Tianjin Sequencing Center & Clinical Lab, China). cDNA synthesis was completed by SMART-Seq^™^ v4 Ultra^™^ Low Input RNA kit for sequencing (Clontech). Samples were sequenced on Illumina machines; paired end, 150bp (PE150) sequencing was performed. RNA-seq data are available at GEO under accession number GSE178979.

### RNA-sequencing analysis

For gene quantification, raw reads were aligned to Reference Genome mm10 in a splice-aware manner using the STAR aligner (43). ENSEMBL gene annotations were quantified using FeatureCounts (44). Differential expression statistics (fold-change and p-value) were computed using edgeR on raw expression counts obtained from quantification. Multi-group analyses were performed to quantify any differences between genotypes using the generalized linear models (GLM), as well as pair-wise tests between sample conditions using exactTest in edgeR (45, 46). rMATS was run with the default settings for the detection of differential alternative splicing events (28).

### Gene ontology analysis

Metascape-Gene Annotation and Analysis Resource was used to perform gene ontology (GO) analysis (47). P-values or q-values were used as representation metrics for ranking enriched terms, as indicated in the Figs..

### Quantitative real-time PCR and splicing validation

Total cellular RNA was isolated with RNeasy Micro Kits with DNase I (Qiagen). cDNA was synthesized using SuperScript^™^ III First-Strand Synthesis System (Invitrogen). RNA levels in LSK+ cells and BM-derived macrophages (BMDMs) were determined by real-time qPCR using SYBR Green master mix (ThermoFisher) (10 min at 95°C, 40 cycles of 15s at 95°C and 1 min at 60°C). Mouse *Gapdh* was used as internal reference and the ΔΔCt method was used to compare expression between samples. For splicing validation, PCR was carried out with Go-Taq Green Master Mix, according to the manufacturer’s protocol (Promega) and products were analyzed on 4% NuSieve 3:1 agarose gels. Primer sequences designed by the NCBI primer-blast tool are shown in *SI Appendix*, Table S1.

### Peripheral blood counts and tissue histology

Peripheral blood was collected and complete blood counts (CBC) were measured (Vet ABC hematology analyzer (scil animal care, Gurnee, IL). Tibiae, spleens and livers were harvested and transferred into 10% neutral buffered formalin for 24h fixation. Tibiae were decalcified in Immunocal overnight, then paraffin-embedded. Longitudinal sections were obtained for tibiae and transverse section for spleen and liver. Five μm sections were stained with hematoxylin and eosin and examined by light microscopy.

### Data and statistical analysis

Statistical analyses were performed by GraphPad Prism. Error bars represent mean ± standard deviation. Tests used to determine significance are indicated in the Fig. legends. Survival curves were constructed by Kaplan-Meier analysis and Log-rank test was used for comparison. No data were excluded from the analysis. All raw data are available at https://data.mendeley.com/datasets/xbr2w4mydr/draft?a=d4a126d2-c3fc-4d30-a408-a20784d5aaa8.

## ACKNOWLEDGEMENTS

We thank David Ryan for help with the mouse breeding and Vemika Chandra, Kristen Lynch, Kostandin Pajcini, Lijian Shao, Wei Tong, and Mark Maienschein-Cline and Pinal Nitish (UIC Research Informatics Core) and Zarema Arbieva (UIC Genomics Core) for advice. Bioinformatics analysis described in the project was performed by the University of Illinois Research Informatics Core, supported in part by the National Center for Advancing Translational Sciences, National Institutes of Health, through Grant UL1TR002003. Supported by NIH Grants R01 AI121275 and R21 AI153594 to SRR.

