## Supplemental Data for "DDX41 is needed for pre-and post-natal hematopoietic stem cell differentiation in mice"

### Figure S1

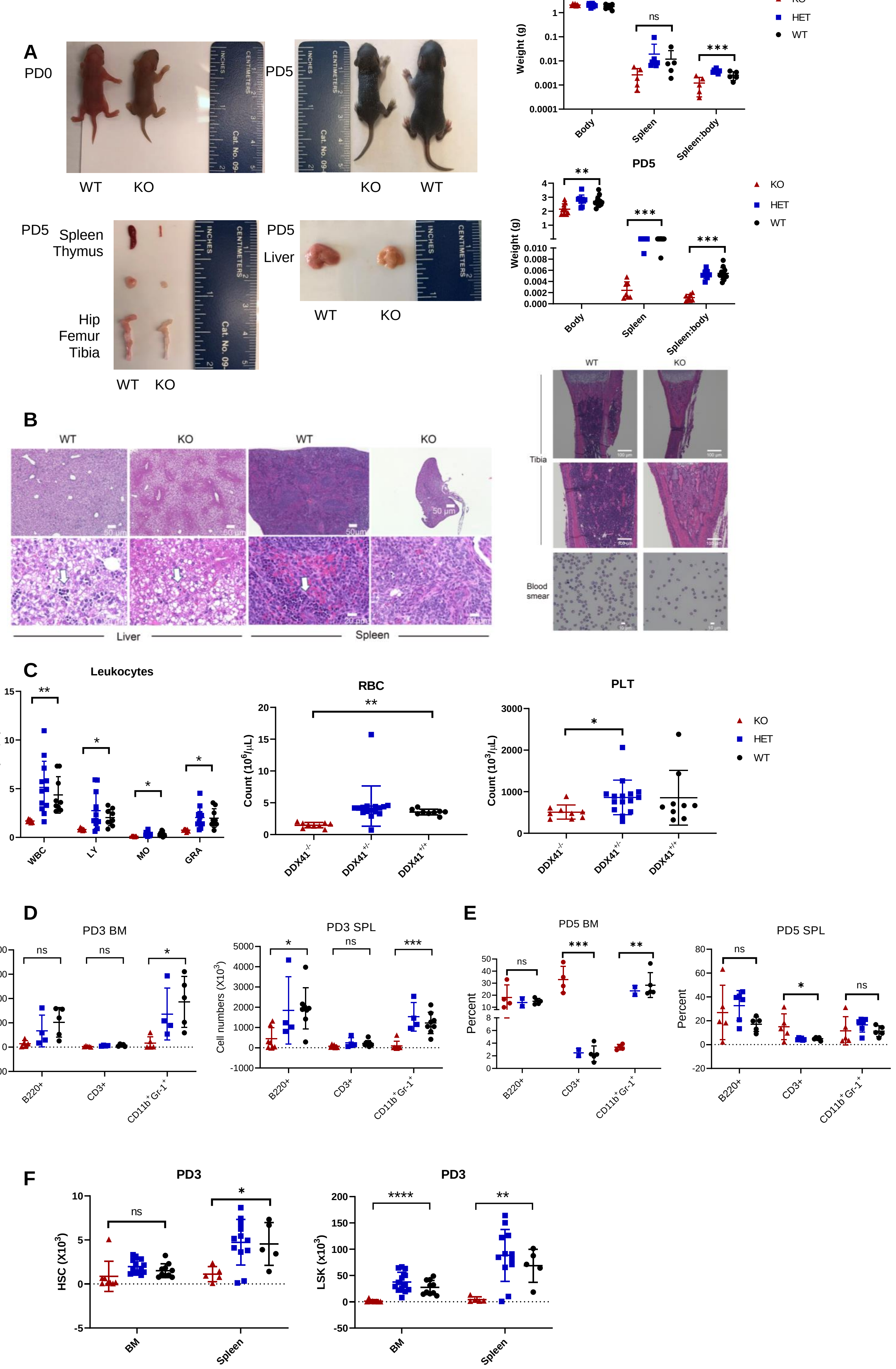
A
PD0
WT KO
PD5
KO WT
PD5 Spleen Thymus
Hip Femur Tibia
WT KO
PD5 Liver
WT KO
PD3
Weight (g)
Body Spleen Spleen:body
*
ns
***
KO HET WT
PD5
**
***
***
B
WT KO WT KO
50 µm
Liver
Spleen
WT KO
Tibia
Blood smear
C
Leukocytes
Count (10^3/µL)
WBC LY MO GRA
RBC
Count (10^6/µL)
DDX41-/- DDX41+/- DDX41+/+
PLT
Count (10^3/µL)
D
PD3 BM
Cell numbers (X10^3)
B220+ CD3+ CD11b+Gr-1+
PD3 SPL
E
PD5 BM
Percent
PD5 SPL
F
PD3
HSC (X10^3)
BM Spleen
LSK (X10^3)
****

**Fig. S1, related to Fig. 1.** DDX41 KO mice fail to thrive. A) DDX41 KO mice appeared normal at birth (PD0) but were runted beginning at PD3-PD5 compared with wt mice. Spleens, liver and bone marrow were all reduced in size, and showed signs of anemia. This was confirmed by body and spleen weights and spleen/body ratio. (PD3: WT mice, n=5; HET mice, n=7; KO mice, n=5; PD5: WT mice, n=12; HET mice, n=9; KO mice, n=7). B) Histological analysis of spleens, thymus, long bones and livers from PD5 KO and WT littermates. Representative hematoxylin and eosin staining of liver, spleen and bone marrow demonstrated anemia and decreased cellularity. Decreased erythroid progenitors in liver and spleen are indicated by white arrows. Wright-Giemsa stain of peripheral blood smears displayed anemia in KO mice. C) CBC of PD3 mice. N ≥ 7 for each group. WBC, white blood cell; LY, lymphocyte; MO, monocyte; GRA, granulocyte. D) HSC cell numbers at PD3 in BM and spleen. E) LSK cells percentage at PD5 in BM and spleen. F) Numbers of HSC and LSK cells at PD3 in BM and spleen. One way ANOVA with post-hoc Tukey's test was used for all comparisons, except PLT in panel C, where unpaired Students T test was used. *, $P \leq 0.05$; **, $P \leq 0.01$; ***, $P \leq 0.001$; ****, $P \leq 0.0001$; ns, not significant.

Numbers of mice

| Panel | Sample | WT | HET | KO |
|---|---|---|---|---|
| C | Leukocytes | 9 | 12 | 7 |
| | RBC/PLT | 9 | 12 | 9 |
| D | BM | 5 | 4 | 5 |
| | SPL | 8 | 4 | 7 |
| E | BM | 5 | 2 | 4 |
| | SPL | 5 | 6 | 5 |
| F | BM | 10 | 13 | 8 |
| | SPL | 5 | 12 | 5 |

#### Figure S2

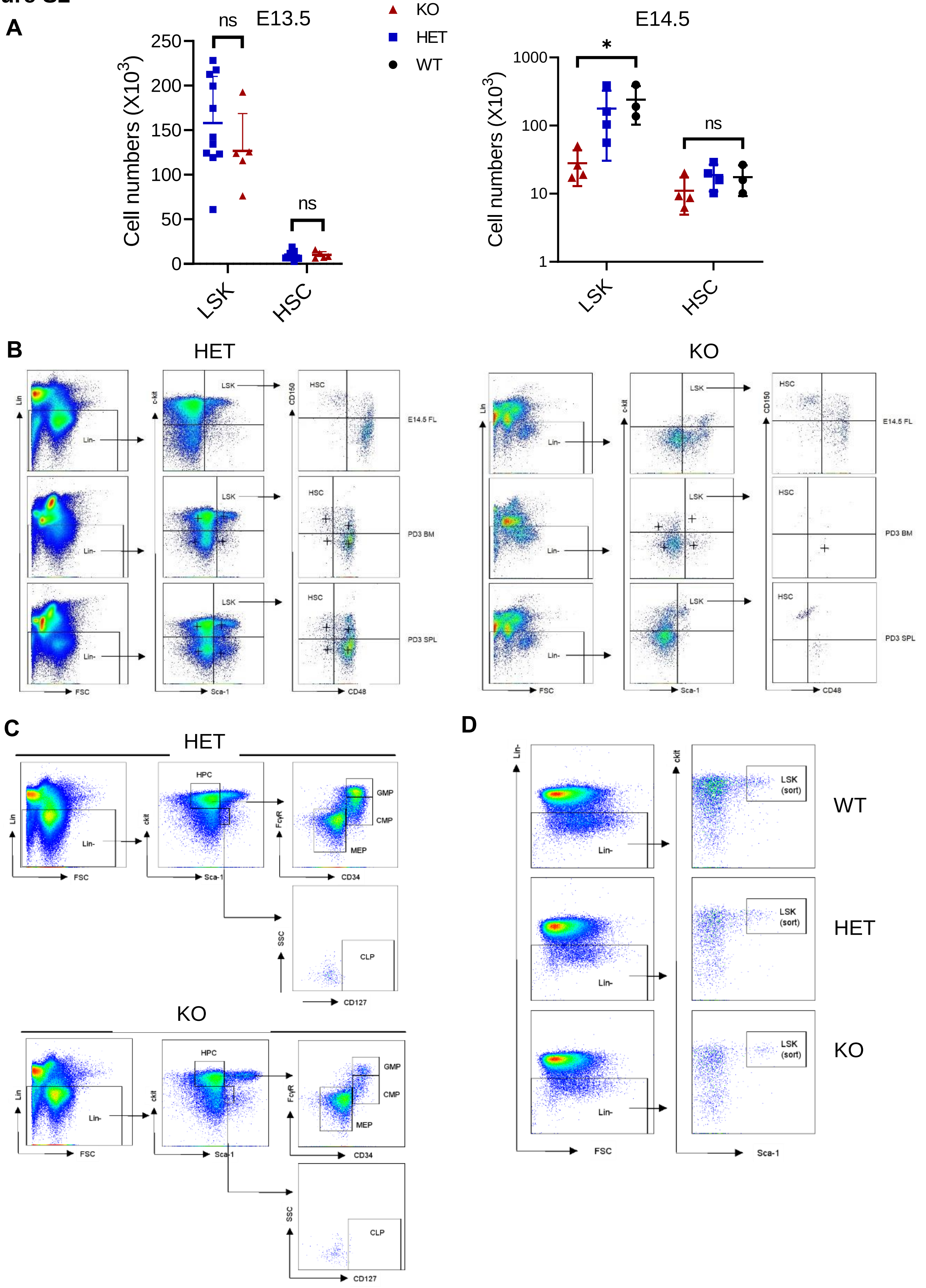

A
E13.5
ns
ns
250
200
150
100
50
0
Cell numbers (X10^3)
LSK
HSC
KO
HET
WT
E14.5
*
ns
1000
100
10
1
Cell numbers (X10^3)
LSK
HSC
B
HET
KO
Lin
Lin-
FSC
c-kit
LSK
Sca-1
CD150
HSC
CD48
E14.5 FL
PD3 BM
PD3 SPL
C
HET
KO
Lin
Lin-
FSC
ckit
HPC
Sca-1
FcγR
GMP
CMP
MEP
CD34
SSC
CLP
CD127
D
Lin-
FSC
ckit
LSK (sort)
Sca-1
WT
HET
KO

**Fig. S2, related to Figs. 1 and 2.** E14.5 HSCs are defective. A) Cells isolated from prenatal liver were examined for hematopoietic stem/progenitor cells (LSK) or hematopoietic stem cells (HSCs) at day E13.5 (unpaired two-tailed t-test) and E14.5 (one-way ANOVA with post-hoc Tukey's test): *, P≤0.05. The mice were derived from 3 litters for each experiment. B) Gating strategy for flow cytometry. Flow cytometry plots showing mouse hematopoietic stem and progenitor cells in HET and KO E14.5 FL, PD3 BM and spleen. Lin-: CD3-/ B220-/ CD11b-/ Gr-1-/ Ter119-, LSK: Lin-/ Sca-1+/ ckit+, HSC: LSK/ CD150+/ CD48-. C) Gating scheme to analyze mouse progenitors in HET and KO E14.5 FLCs. HPC: Lin-/ ckit+, GMP: HPC/ CD34+/ FcγR+, CMP: HPC/ CD34+/ FcγR-, MEP: HPC/ CD34-/ FcγR-, CLP: Lin-/ Sca-1low/ citlow/ CD127+. D) Gating scheme for sorting of LSK cells from WT, HET and KO E14.5 FL.

Numbers of mice

| Sample | WT | HET | KO |
|---|---|---|---|
| E13.5 | 3 | 4 | 4 |
| E14.5 | - | 11 | 5 |

#### Figure S3

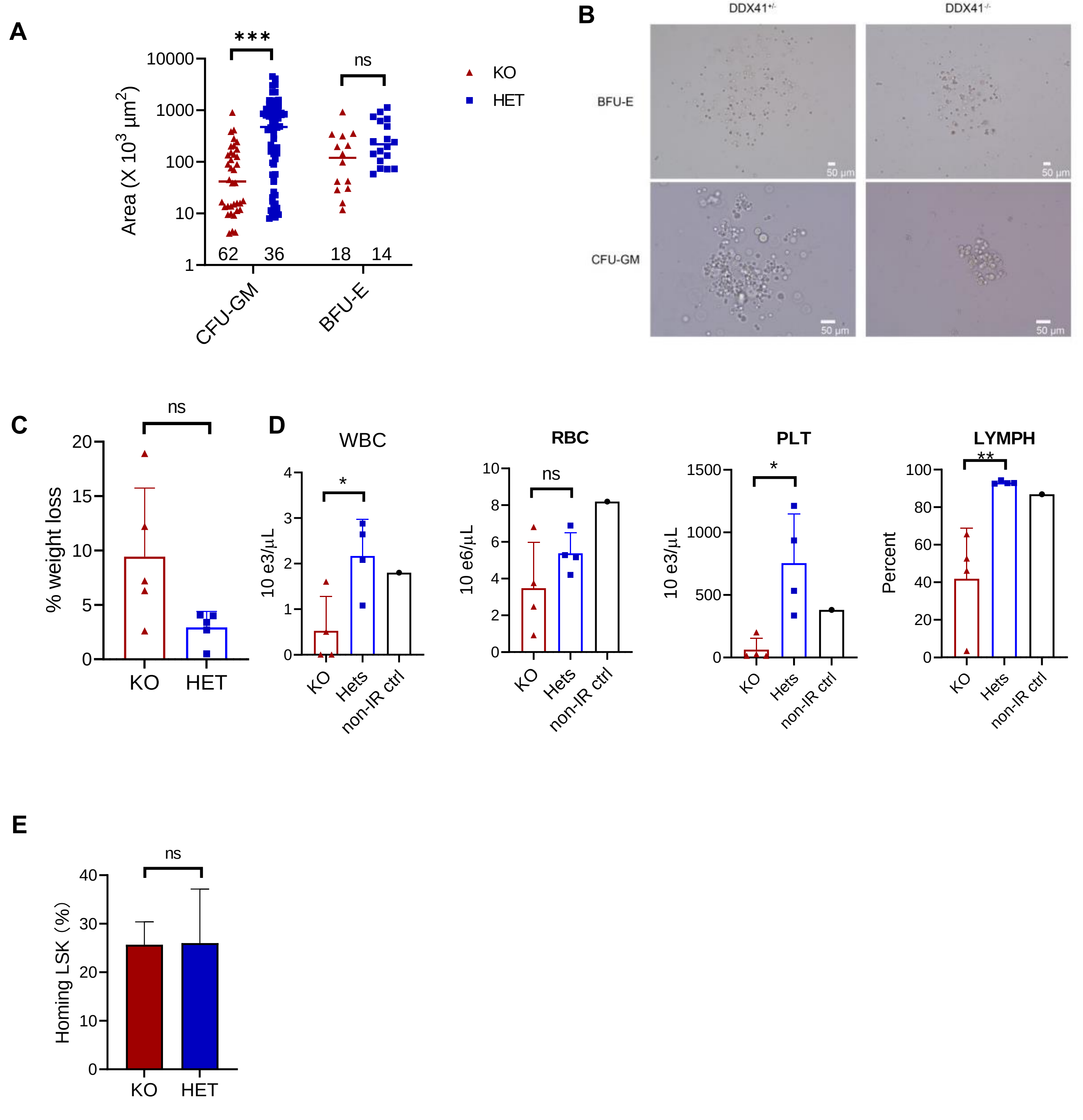

A
***
ns
10000
1000
100
10
1
Area (X 10³ µm²)
62
36
18
14
CFU-GM
BFU-E
KO
HET
B
DDX41+/-
DDX41-/-
BFU-E
CFU-GM
50 µm
C
ns
% weight loss
KO
HET
D
WBC
RBC
PLT
LYMPH
10 e3/µL
10 e6/µL
Percent
*
**
KO
Hets
non-IR ctrl
E
Homing LSK (%)

**Fig. S3, related to Fig. 3.** Lack of DDX41 in HSCs affects their differentiation capacity. A) Colony size was measured in CFU-assay for CFU-GM and BFU-E. BZ-X Analyzing software was used for measurement. Shown are the averages from three independent experiments ± STD. Numbers of colonies are shown above the x axis. B) Representative capture of CFU-GM and BFU-E from hets and KO E14.5 FLCs under the same magnification. The scale bar shows 50μm. C) Weight loss in recipients 2 weeks after BM transplantation. N= 5 for each group. D) CBC test of the BM transplantation recipients. N=4 for each group. N=1 for non-ablated control. E) Homing of CD45.1+ E14.5 FL LSK cells (CD45.2) in the BM of recipients (CD45.1) at 16 h after injection. N=3 for each group. Unpaired two-tailed t-tests were used to determine significance. *, $P \leq 0.05$; **, $P \leq 0.01$; ***, $P \leq 0.0001$.

#### Figure S4

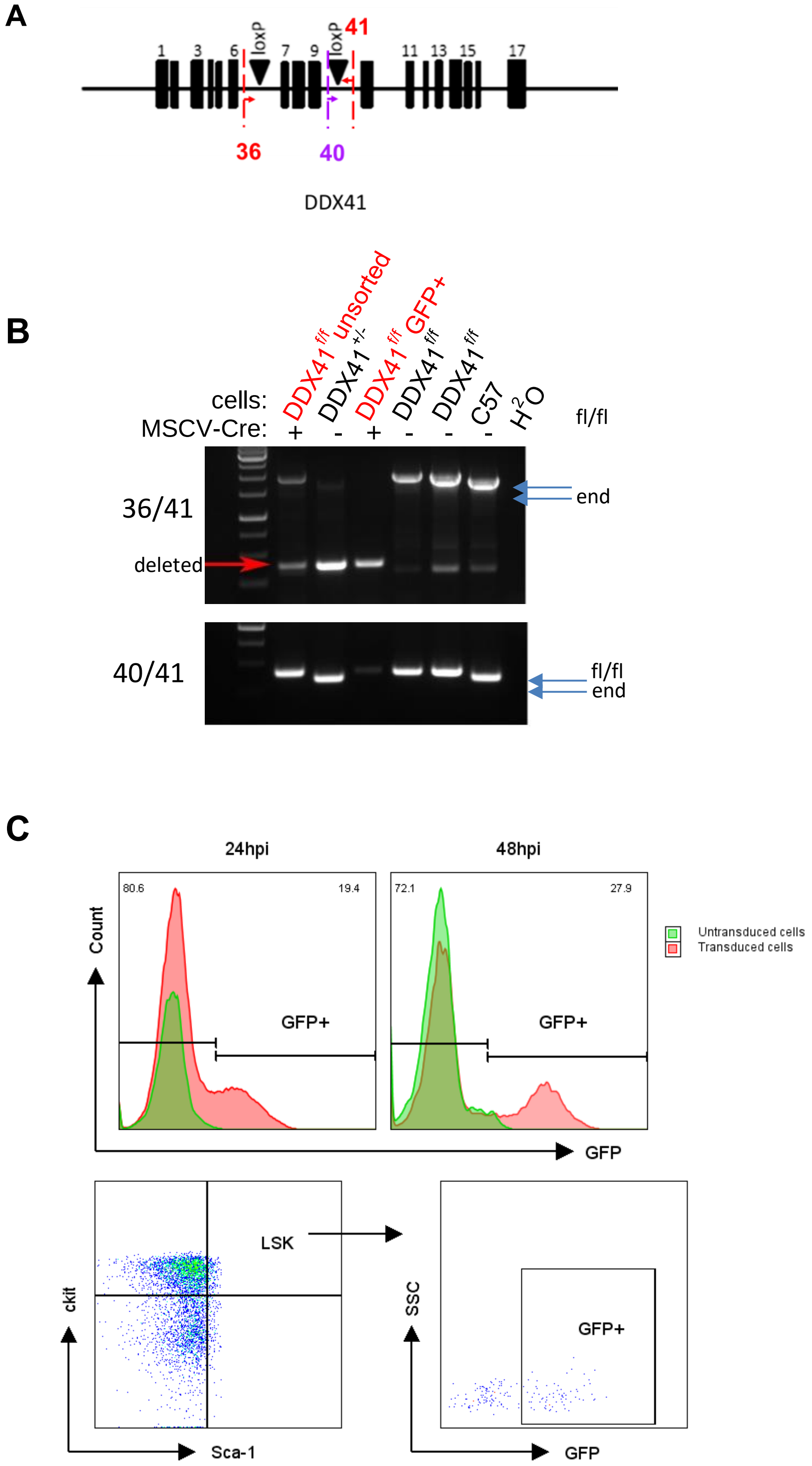

A
1
3
6
loxP
7
9
loxP
41
11
13
15
17
36
40
DDX41
B
cells:
MSCV-Cre:
DDX41 f/f unsorted
DDX41 +/-
DDX41 f/f GFP+
DDX41 f/f
DDX41 f/f
C57
H2O
+
-
+
-
-
-
fl/fl
36/41
deleted
end
40/41
fl/fl
end
C
24hpi
48hpi
80.6
19.4
72.1
27.9
Count
GFP+
GFP+
GFP
Untransduced cells
Transduced cells
LSK
ckit
Sca-1
SSC
GFP+
GFP

**Fig. S4, related to Fig. 3.** DDX41-deficient BM failed to expand in vivo. A) Diagram of murine *Ddx41* with loxP sites. Arrows indicate sets of primers used for genotyping. B) PCR showing deletion of exons 7-9 in Cre-transduced cells. Red arrow indicates the product after gene deletion. Shown is the PCR before (unsorted) and after sorting for GFP expression (GFP+). C) LSK population and GFP+ cells were analyzed by flow cytometry at 24 and 48 h after transduction.

#### Figure S5

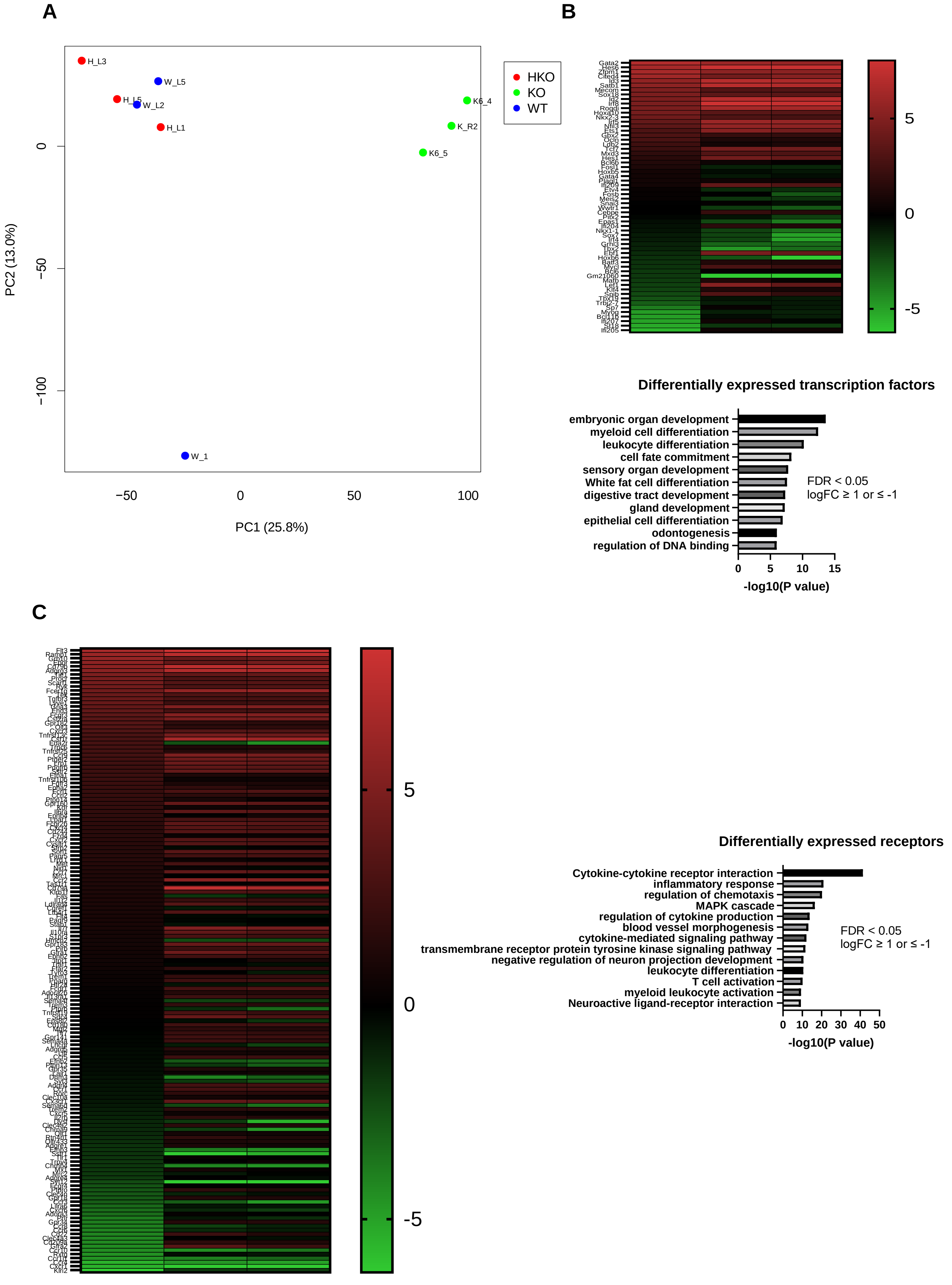

A
B
C
H_L3
W_L5
H_L5
W_L2
H_L1
W_1
K6_4
K_R2
K6_5
HKO
KO
WT
PC1 (25.8%)
PC2 (13.0%)
−50
0
50
100
−100
Differentially expressed transcription factors
embryonic organ development
myeloid cell differentiation
leukocyte differentiation
cell fate commitment
sensory organ development
White fat cell differentiation
digestive tract development
gland development
epithelial cell differentiation
odontogenesis
regulation of DNA binding
FDR < 0.05
logFC ≥ 1 or ≤ -1
-log10(P value)
Differentially expressed receptors
Cytokine-cytokine receptor interaction
inflammatory response
regulation of chemotaxis
MAPK cascade
regulation of cytokine production
blood vessel morphogenesis
cytokine-mediated signaling pathway
transmembrane receptor protein tyrosine kinase signaling pathway
negative regulation of neuron projection development
leukocyte differentiation
T cell activation
myeloid leukocyte activation
Neuroactive ligand-receptor interaction
FDR < 0.05
logFC ≥ 1 or ≤ -1
-log10(P value)

**Fig. S5, related to Fig. 4.** Distinct gene expression patterns in KO and WT/het cells. A. Principal component analysis (PCA) plot showing distinct clusters between KO and WT/het cells in gene expression data. B. A heatmap (upper) representation of differentially expressed transcription factors (TF) between KO, WT and het. N=3 for each group. Gene oncology (GO) enrichment analysis (lower) of differential expressed TFs. logFC (KO/WT) ≥ 1 or ≤ -1, FDR < 0.05. C. A heatmap representation (left) and GO analysis (right) of differentially expressed receptors.

**Figure S6**

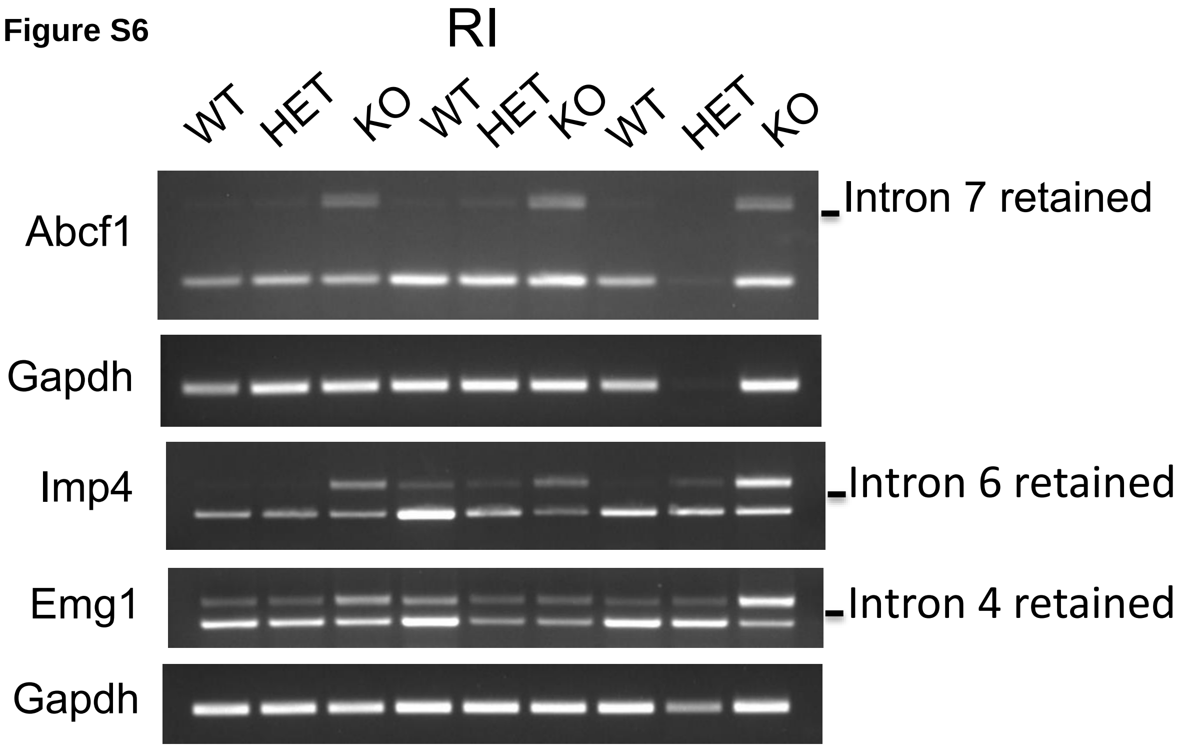

| Gene | Function | RE effect | Inclusion Difference | |
|---|---|---|---|---|
| | | | WT-KO | HET-KO |
| Abcf1 | ATP binding cassette transporter | ? | -0.196 | -0.2 |
| Imp4 | U3 snoRNP complex | Premature termination | -0.332 | -0.333 |
| Emg1 | rRNA Ψ U methyltransferase | Premature termination | -0.212 | -0.211 |

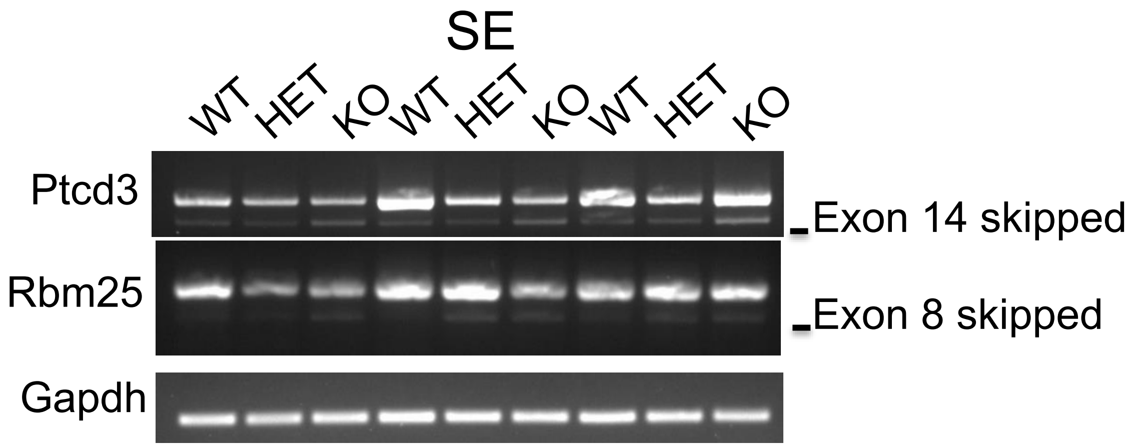

| Gene | Function | SE effect | Inclusion Difference | |
|---|---|---|---|---|
| | | | WT-KO | HET-KO |
| Ptcd3 | Mitochondrial RNA binding protein | RNA binding | 0.149 | 0.169 |
| Rbm25 | RNA-binding protein, regulator of alternative pre-mRNA splicing | ? | 0.1 | 0.098 |

**Fig. S6, related to Fig. 5.** DDX41 regulates splicing in LSK cells. PCR validation of retained intron in Abcf1, Imp4, Emg1 (upper). PCR validation of skipped exon in Ptcd3 and Rbm25 (lower). Tables below the PCR gels summarize gene function and inclusion difference in WT vs. KO and HET vs. KO. Abbreviations: NMD, nonsense-mediated decay.

#### Figure S7

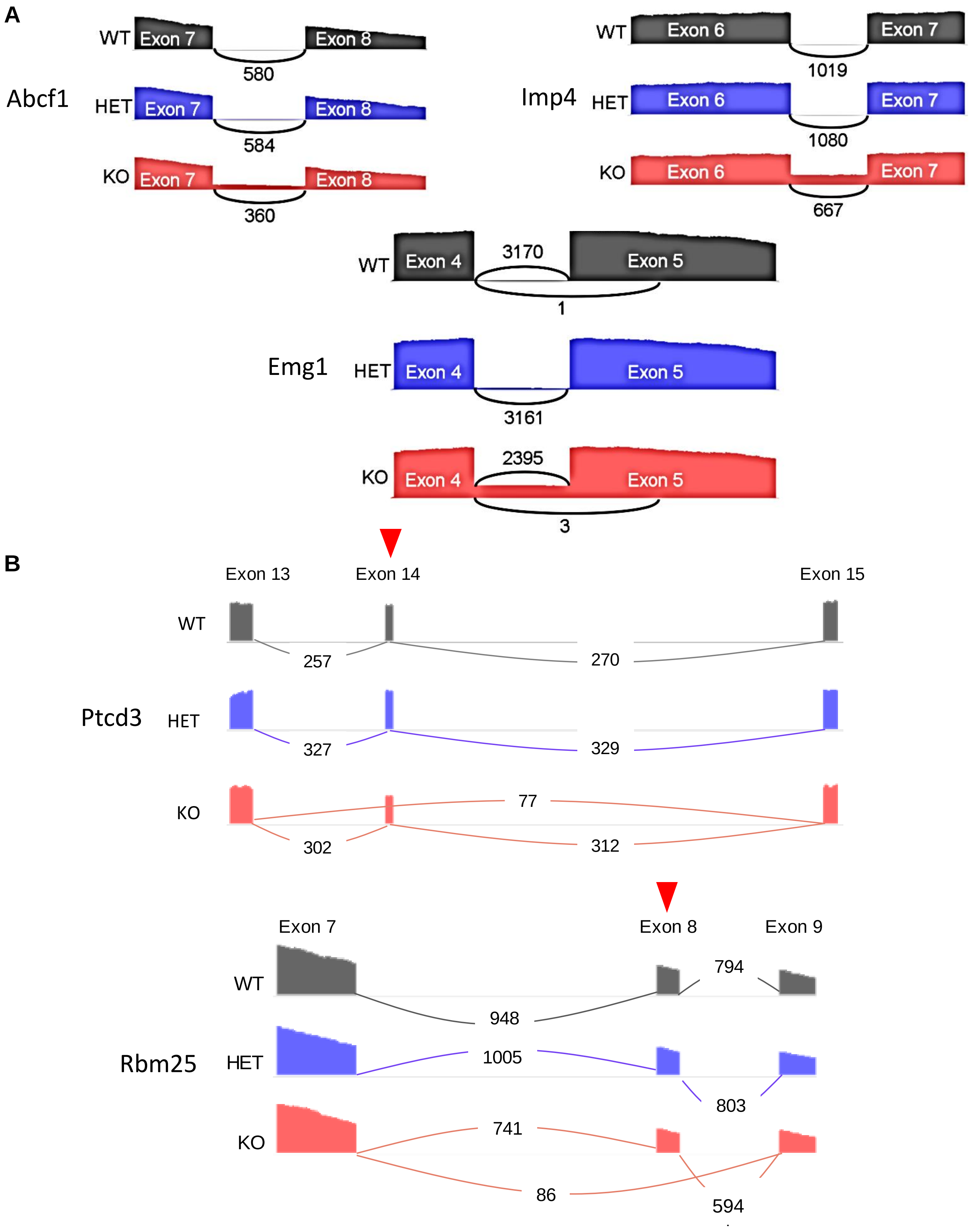
A
Abcf1
WT
Exon 7
Exon 8
580
HET
Exon 7
Exon 8
584
KO
Exon 7
Exon 8
360
Imp4
WT
Exon 6
Exon 7
1019
HET
Exon 6
Exon 7
1080
KO
Exon 6
Exon 7
667
Emg1
WT
Exon 4
3170
Exon 5
1
HET
Exon 4
Exon 5
3161
KO
Exon 4
2395
Exon 5
3
B
Ptcd3
Exon 13
Exon 14
Exon 15
WT
257
270
HET
327
329
KO
77
302
312
Rbm25
Exon 7
Exon 8
Exon 9
WT
794
948
HET
1005
803
KO
741
86
594

**Fig S7, related to Fig. 5**. Differential splicing between WT, HET and KO cells. A) Sashimi plots representing retained introns in KO cells for Abcf1, Imp4 and Emg1 are shown. Numbers indicate reads from RNA-seq. B) Sashimi plots representing skipped exons for Ptcd3 and Rbm25 in KO cells are shown. Red triangles stand for skipped exons.

Figure S8

A

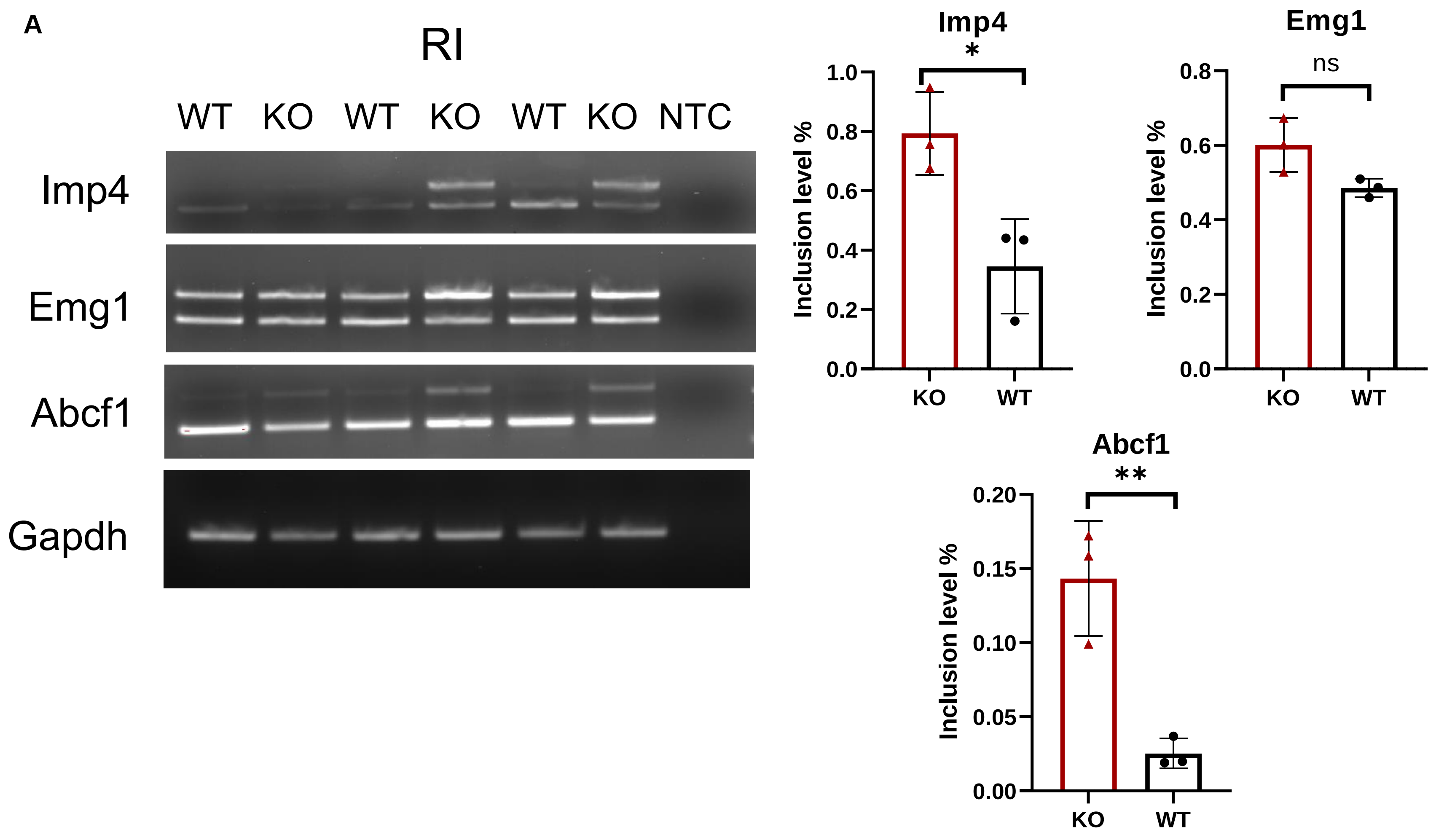
RI
WT KO WT KO WT KO NTC
Imp4
Emg1
Abcf1
Gapdh
Imp4
*
Inclusion level %
1.0
0.8
0.6
0.4
0.2
0.0
KO
WT
Emg1
ns
0.8
0.6
0.4
0.2
0.0
Abcf1
**
0.20
0.15
0.10
0.05
0.00

B

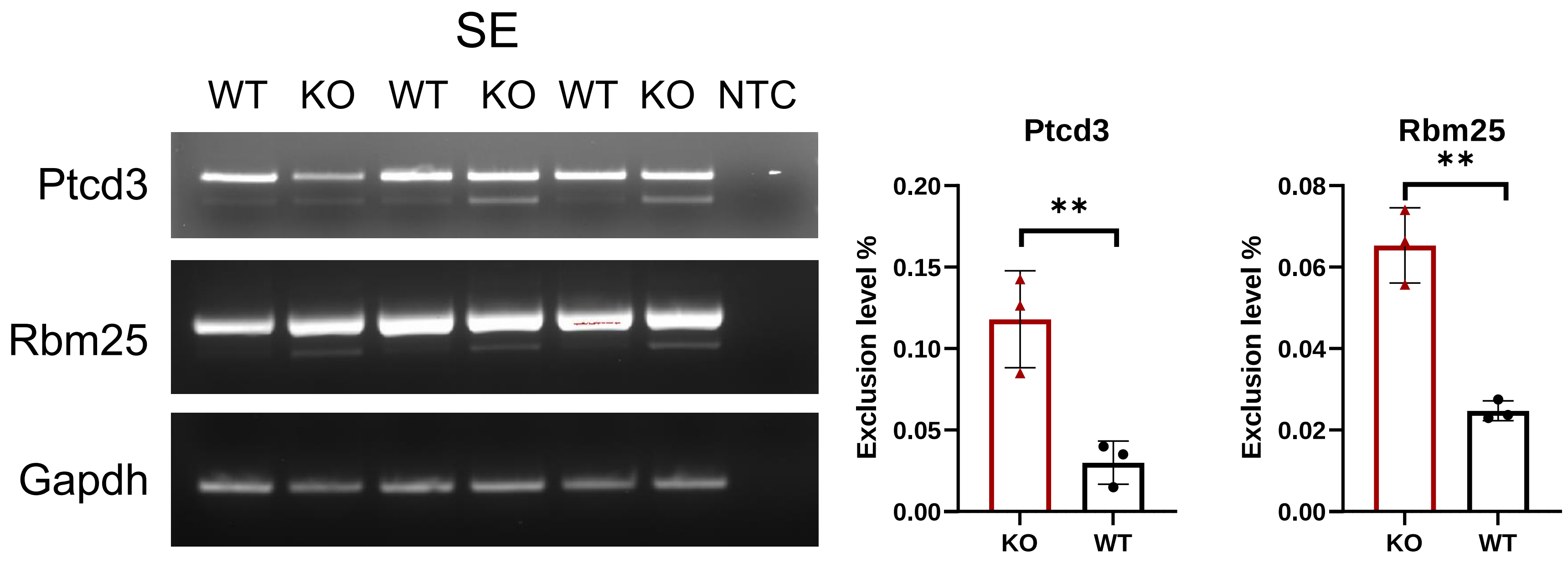
SE
WT KO WT KO WT KO NTC
Ptcd3
Rbm25
Gapdh
Ptcd3
**
Exclusion level %
0.20
0.15
0.10
0.05
0.00
KO
WT
Rbm25
**
0.08
0.06
0.04
0.02
0.00

**Fig S8, related to Fig. 5.** DDX41 regulates splicing in BMDMs. A) Exon skipping of Ptcd3 and Rbm25 were analyzed by PCR (left). PCR products were quantified, and the exclusion level is indicated (right). B) Intron retention of Imp4, Emg1, Abcf1 was analyzed by PCR (left). The inclusion level is shown (right). The inclusion or exclusion levels were calculated as described in Fig. 5. Unpaired two-tailed t-test was used to determine significance: *, $P \leq 0.05$, **, $P \leq 0.01$.

**Table S1.** Antibody (for flow cytometry and FACS sorting) and reagents.

| Marker | Manufacturer | Catalogue number | Dilution | Fluorescence | Clone |
|---|---|---|---|---|---|
| CD3 | ThermoFisher, Carlsbad, CA | 25-0031-81 | 1:200 | PE/Cyanine7 | 145-2C11 |
| CD3 | SouthernBiotech, Birmingham, AL | 1530-02 | 1:400 | FITC | 145-2C11 |
| CD11b | Biolegend, San Diego, CA | 101206 | 1:500 | FITC | M1/70 |
| CD11b | Biolegend, San Diego, CA | 101227 | 1:500 | PerCP/Cyanine5.5 | M1/70 |
| Gr-1 | Biolegend, San Diego, CA | 108408 | 1:500 | PE | RB6-8C5 |
| Gr-1 | Biolegend, San Diego, CA | 108437 | 1:500 | Brilliant Violet 510™ | RB6-8C5 |
| Gr-1 | Biolegend, San Diego, CA | 108406 | 1:500 | FITC | RB6-8C5 |
| B220 | Biolegend, San Diego, CA | 103212 | 1:300 | APC | RA3-6B2 |
| B220 | Biolegend, San Diego, CA | 103257 | 1:200 | PE/Dazzle™594 | RA3-6B2 |
| B220 | Biolegend, San Diego, CA | 103206 | 1:200 | FITC | RA3-6B2 |
| CD11b | Biolegend, San Diego, CA | 101206 | 1:400 | FITC | M1/70 |
| ckit | Biolegend, San Diego, CA | 135108 | 1:200 | APC | ACK2 |
| Sca-1 | Biolegend, San Diego, CA | 122508 | 1:400 | PE | E13-161.7 |
| CD150 | Biolegend, San Diego, CA | 115913 | 1:300 | PE/Cy7 | TC15-12F12.2 |
| CD48 | Biolegend, San Diego, CA | 103445 | 1:600 | APC/FireTM750 | HM48-1 |

| | | | | | |
|---|---|---|---|---|---|
| CD45.1 | Biolegend, San Diego, CA | 110724 | 1:200 | AF700 | A20 |
| CD45.1 | BD Biosciences, San Jose, CA | 561872 | 1:200 | PE | A20 |
| CD45.2 | BD Biosciences, San Jose, CA | 560694 | 1:200 | APC/Cy7 | 104 |
| CD16/32 | Biolegend, San Diego, CA | 156611 | 1:500 | APC/Cyanine7 | S17011E |
| CD34 | Biolegend, San Diego, CA | 128617 | 1:200 | PE/Cy7 | HM34 |
| CD127 | ThermoFisher, Carlsbad, CA | 45-1271-80 | 1:300 | PerCP/Cy5.5 | A7R34 |
| Ter119 | Biolegend, San Diego, CA | 116223 | 1:400 | APC/Cyanine7 | TER-119 |
| Ter119 | Biolegend, San Diego, CA | 116206 | 1:300 | FITC | TER-119 |
| Murine IL-6 | Peprotech,Cranbury, NJ | 216-16 | | | |
| Murine IL-3 | Peprotech,Cranbury, NJ | 213-13 | | | |
| Murine SCF | Peprotech,Cranbury, NJ | 250-03 | | | |
| Murine TPO | Peprotech,Cranbury, NJ | 315-14 | | | |

**Table S2.** Primer sequences for genotyping and RT PCR to validate splicing analysis.

| | **Forward** | **Reverse** | **Experiments** |
|---|---|---|---|
| VavCre | GCGGTCTGGCAGTAAAAACTATC | GTGAAACAGCATTGCTGTCACTT | Genotyping |
| WT allele | CCAGTAGCACCTCTGCTTGG | CTGCCTCTCAGCACTTCCC | Genotyping |
| KO allele | GGAGGCATAACCACCACA | CTGCCTCTCAGCACTTCCC | Genotyping |
| **Gene name** | **Forward** | **Reverse** | **Experiments** |
| Ptcd3 | CATCATCAAGGTCCTGTGGAG | GGTTGCACAGAAAGTGAAACCA | RT-PCR |
| Rbm25 | AACGCCAGACCAGAAACTGT | CTGAATTTGCTGATCTCTCGGG | RT-PCR |
| Abcf1 | TGTCCTTCCCTTGCTGAGGAG | GGCTCAGGAGTAAAAAGGGAAAG | RT-PCR |
| Imp4 | ATCTGCCCTTCGGTCCTACT | TTGCAAAGGTGATGACCCGA | RT-PCR |
| Emg1 | CAACTGGGTCACTACTGGGC | ACACAAACTGAGCGTCCGAG | RT-PCR |
| Vpreb2 | ACACTCAGACAAGCACCAGG | CCCCACAGCGCAGTAATACA | RT-PCR |
| Clec7a | CTTCAGCACTCAAGACATCCATAA | CAAGGTGAAGATGGAGCCTG | RT-PCR |
| Eda2r | ACCAAGAATGCATCCCATGTA | CCTAGGGGGAACAGTGTGTG | RT-PCR |
| Ddx41 | GTCGGGACATGATCGGCATT | CCCCTCTCGCTTGGAGAAAG | RT-PCR |
| Gapdh | GGAGAGTGTTTCCTCGTCCC | ATGAAGGGGTCGTTGATGGC | RT-PCR |

**Dataset S1.** Differentially expressed genes in KO vs. WT, Qvalue < 0.05, log2FC ≥1 or ≤ -1. Related to Figure 4.

**Dataset S2.** Differential RI events in WT v.s KO. FDR < 0.05, |Inc level difference| ≥ 0.1. Related to Figure 5.

**Dataset S3.** Differential SE events in WT v.s KO. FDR < 0.05, |Inc level difference| ≥ 0.1. Related to Figure 5.

**Dataset S4.** Differential A3SS events in WT v.s KO. FDR < 0.05, |Inc level difference| ≥ 0.1. Related to Figure 5.

**Dataset S5.** Differential A5SS events in WT v.s KO. FDR < 0.05, |Inc level difference| ≥ 0.1. Related to Figure 5.

**Dataset S6.** Differential MXE events in WT v.s KO. FDR < 0.05, |Inc level difference| ≥ 0.1. Related to Figure 5.

**Dataset S7.** Differential spliced events in WT v.s KO. FDR < 0.05, |Inc level difference| ≥ 0.1.
